# Differential genetic resistance identified in *Parastagonospora nodorum* and *Pyrenophora tritici-repentis*-wheat pathosystems

**DOI:** 10.64898/2026.06.12.731808

**Authors:** Huyen T. T. Phan, Eiko Furuki, Fiona Kamphuis, Kasia Rybak, Leon Lenzo, Catherine F. Cupitt, Kalai Marathamuthu, Pao Theen See

## Abstract

Septoria nodorum blotch (SNB) and tan spot (TS) wheat diseases are caused by necrotrophic fungal pathogens *Parastagonospora nodorum* (Pn) and *Pyrenophora tritici-repentis* (Ptr), respectively. Although recognised as premier model pathosystems for our understanding of necrotrophic effectors, no resistance mechanism has been reported in both diseases. Here, two SNB and TS resistance wheat lines (‘56:ZWB11’ and ‘105:ZIF14’) derived from the Australian national germplasm evaluation programme (CAIGE) were used to develop a double haploid mapping population. Two Pn and Ptr isolates of different pathotypes, their respective culture filtrates and effector SnTox267 were evaluated on the population. Genetic analysis of Ptr conidial inoculation of race 1 and race 2 identified a major resistance quantitative trait locus (QTL) (*QTs.cur*–1B) on chromosome 1B, while resistance to SNB was explained by several minor QTL. SnTox267 sensitivity was mapped to six locations (2A2, 2A3, 2B1, 2D3, 5B and 7B1) with only one QTL co-localized to known corresponding gene *Snn7*. Sensitivity loci 5B and 7B1 also conferred SNB resistance at seedling and adult stages. Two QTL on chromosome 2D1 and 7B2 were common in both SNB and TS, associated with disease at seedling stage and culture filtrate bioactivity, respectively. Resistance responses of ‘56:ZWB11’ and ‘105:ZIF14’ were confirmed cytologically, however, distinct responses were observed on wounded leaves. The defence responses were more effective against Ptr, while resistance to Pn infection was likely a combination of lack of susceptibility and effective physical barriers. Overall results demonstrated the distinction between the underlying resistance mechanisms to TS and SNB.

## Introduction

Bread and durum wheats (*Triticum aestivum* L. and *Triticum turgidum* spp*. durum*) are among the most important crops worldwide, accounting for around 20% of global human calorie intake (Savary et al. 2019). However, wheat production is often limited primarily by diseases. Fungal wheat diseases are leading causes of global yield losses with major threats including tan spot (TS, also known as yellow spot) and septoria nodorum blotch (SNB, also known as septoria glume blotch when infection occurs on wheat heads). SNB and TS are caused by necrotrophic fungi *Parastagonospora nodorum* (Pn) and *Pyrenophora tritici-repentis* (Ptr), respectively. Losses in wheat yield associated with these diseases typically ranged from 5% to 31% with up to 60% reported in highly susceptible cultivars (Bhathal et al. 2003; Cook et al. 2024; Ficke et al. 2018; Rees and Platz 1989). The losses were due to a reduction in photosynthetic area of leaves, which affects grain quantity and quality, including low weight and shrivelled grains, reduced number of kernels per head (De Wolf et al. 1998; Ficke et al. 2018; Shabeer and Bockus 1988) and grain discolouration (Fernandez et al. 1998; Ficke et al. 2018).

SNB and TS disease incidences are driven by several interactions between necrotrophic effectors (NEs) produced by the pathogens and corresponding dominant susceptibility genes of the hosts. The interactions follow an inverse gene-for-gene manner where recognition of NE by the host will result in an effector-triggered response that leads to disease susceptibility (reviewed by Faris et al. 2013; Friesen and Faris 2010; Liu et al. 2017; Peters Haugrud et al. 2019). To date, nine of such interactions (ToxA*-Tsn1,* SnTox1*-Snn1*, SnTox3*-Snn3_B,* SnTox3*-Snn3_D,* Tox5*-Snn5 and* Tox267*-Snn2/6/7*) have been described in details in the Pn-wheat pathosystem, while in Ptr-wheat pathosystem, there are three interactions (ToxA*-Tsn1,* ToxC*-Tsc1,* ToxB*-*Tsc2) (Abeysekara et al. 2012; Abeysekara et al. 2009; Abeysekara et al. 2010; Ciuffetti et al. 2010; Effertz et al. 2002; Faris et al. 2010b; Friesen and Faris 2004; Friesen et al. 2008; Gao et al. 2015; Kariyawasam et al. 2022; Liu et al. 2006a; Liu et al. 2012; Manning and Ciuffetti 2005; Richards et al. 2022; Shi et al. 2016; Zhang et al. 2011). Of the multiple effector-host gene interactions, ToxA -*Tsn1* interaction is common in both SNB and TS pathosystems and plays a significant role in both diseases. In fact, genomic analysis indicated that ToxA could have been introduced to Ptr through interspecific horizontal gene transfer from Pn (Friesen et al. 2006; McDonald et al. 2019).

Thus far, Ptr is classified into eight races based on their possession of the three effectors (ToxA, ToxB and ToxC) and the differential disease responses on a set of wheat lines harbouring the respective corresponding genes (*Tsn1*, *Tsc2* and *Tsc1*) (Lamari et al. 1995; Lamari et al. 2003). These races induce different virulence profiles from no symptom to tan necrosis and/or extensive chlorosis depending on the susceptibility of the host, with ToxA-producing isolates inducing necrosis on *Tsn1* line while ToxB or ToxC-producing isolates induce chlorosis on *Tsc2* and *Tsc1*, respectively. The combination of the different effector-host interactions define the eight races (Race 1 – ToxA and ToxC, Race 2 – ToxA only, Race 3 – ToxC only, Race 4 – no effector, Race 5 – ToxB only, Race 6 – ToxB and ToxC, Race 7 – ToxA and ToxB and Race 8 – ToxA, ToxB and ToxC); however, recent studies have challenged the traditional race classification of Ptr with some isolates found to not conform to any of the race categories (Abdullah et al. 2017; Ali et al. 2010; Moolhuijzen et al. 2022). Unlike Ptr, virulence profiles of Pn isolates are postulated to be more complex. Despite SNB disease being arguably one of the most well-studied pathosystems in NE biology, there is no universal pathotype classification or races defined to date.

Host resistance/susceptibility to Ptr ranges from qualitative (Anderson et al. 1999; Faris et al. 1996; Singh and Hughes 2006; Tadesse et al. 2007) to quantitative genetics (Cheong et al. 2004; Faris and Friesen 2005; Friesen and Faris 2004; Singh et al. 2008). Liu et al. (2020) mega-QTL analysis studies reported over 100 QTL associated with resistance/susceptibility to TS. The QTL were identified from various isolates of different races, locations and environments, thus suggesting that the Ptr-wheat system is more complex than the standard race classification system would suggest. These QTL, however, can be grouped into 19 core regions, which include the three known NE sensitivity QTL located on chromosome 5B (*Tsn1*), 1A (*Tsc1*) and 2B (*Tsc2*).

For the Pn-wheat pathosystem, over 200 QTL were reported for SNB and septoria glume blotch (reviewed by Downie et al. 2021). Although there are studies that have identified a single dominant gene conferring resistance/susceptibility (Ma and Hughes 1995; Murphy et al. 2000), most showed a quantitative nature of SNB resistance at adult and seedling stages in both foliar and glume diseases (Adhikari et al. 2011; Czembor et al. 2019; Downie et al. 2018; Francki et al. 2020; Hu et al. 2019; Lin et al. 2020; Phan et al. 2021; Ruud et al. 2018; Wicki et al. 1999). SNB related QTL have been ide in almost all 21 wheat chromosomes from across various studies; however, there are ‘hot spots” where multiple QTL detected from different geographical isolates, environments and growth conditions co-locate on chromosomes 1A, 1B, 2A, 2D, 3A, 3B, 4B, 5A, 5B, 6A, 7A, 7B and 7D (reviewed by Downie et al. 2021). Similar to Ptr-wheat pathosystem, host sensitivity to Pn NE ToxA, SnTox1, SnTox3, SnTox5 and SnTox267 also mapped to the corresponding QTL located on 5BL, 1BS, 5BS, 4BL and 2DS/6AS/2DL, respectively, all with significant contributions to SNB disease development (Abeysekara et al. 2012; Abeysekara et al. 2009; Faris et al. 2010a; Friesen et al. 2012; Friesen et al. 2008; Gao et al. 2015; Kariyawasam et al. 2022; Liu et al. 2006b; Liu et al. 2012; Phan et al. 2016; Richards et al. 2022; Zhang et al. 2011).

Currently, SNB and TS disease management practices include fungicide applications and crop rotation; however, the most effective, economical and environmentally sound approach for controlling these fungal diseases is breeding resistant varieties for cultivation. In Australia, a national program referred to as CAIGE, the CIMMYT (International Maize and Wheat Improvement Centre) Australia ICARDA (International Centre for Agricultural Research in the Dry Areas) Germplasm Evaluation, was initiated in 2007 to streamline the importation, quarantine, seed-multiplication and traits assessment processes of wheat germplasm developed by the Consultative Group on International Agricultural Research (CGIAR), for Australian wheat breeders and researchers (Trethowan et al. 2024). Through this program, approximately 250 wheat lines of diverse genetic materials are made available to the industry annually for selection of beneficial alleles/traits in breeding programs. In this study, we evaluated and identified SNB and TS resistant wheat lines from the CAIGE germplasm and characterised the underlying genetic resistance of these novel resources in both diseases. Information obtained from this study will help to facilitate breeding for SNB and TS resistance in Australian wheat cultivars.

## Material and Methods

### Biological materials

Information of wheat lines from CAIGE collection that displayed SNB and TS resistance in Australian field evaluations (https://www.caigeproject.org.au/germplasm-evaluation/bread/disease-screening-2/) is provided in Supplementary Table S1. Six to eight Australian commercial wheat cultivars (Supplementary Table S2) and ‘Chinese Spring’ were used as controls. For SNB disease assessments, a panel of 12 genetically diverse Australian Pn isolates (Supplementary Table S3) identified from previous study (Phan et al. 2020) were used for the initial disease assessment and two isolates, Northam_WGT and 16FG168 with different pathogenicity profiles (Jones et al. 2024) were used in sequential assessments; meanwhile two Ptr isolates (W0238WY2 and W0224MG) that represent the Australian Ptr races, race1 and race 2, respectively (See et al. 2023) were selected for TS disease assessments..

To analyse the resistance mechanism, a double haploid wheat population designated as DH105 x 56 was developed from a cross between the CAIGE resistant lines ‘56:ZWB11’ and ‘105:ZIF14’. Plant bioassays using purified effector proteins were conducted to screen for SnTox3 and SnTox267 sensitivities as described under methodology section ‘Effectors and crude culture filtrates bioassays’. No further screening for ToxA, ToxB, SnTox1 and SnTox5 was conducted since both lines were insensitive to these effectors. Excluding the aforementioned NE-host gene interactions reduced the complexity of the QTL analysis.

### Comparison of means

ANOVA analysis was used to compare the means of SNB and TS disease scores using R v4.5.2 (R Core Team 2021). Disease severity ranking among wheat lines was carried out using agricolae v1.4.0 (Mendiburu and Yaseen 2020). Statistical significance for all comparisons was defined as *p* < 0.05.

### Plant infection assay

Plant infection assays at seedling stage was carried out in controlled growth environment as described by Phan et al. (2016) for Pn infection and See et al. (2023) for Ptr. All 241 DH lines were planted irandomised design with three replicates. For the infections, pots (10 cm in diameter) containing vermiculite were sown with six seeds and grown at 22 ± 1 °C under a 12-h day/night cycle. Pots were fertilised with Thrive water-soluble fertilizer (Yates, New South Wales, Australia) according to the manufacturer’s recommended application rate. Depending on the germination, four to six seedlings per line in a pot were considered as one biological replicate. Two-week old seedlings were sprayed with spore inoculum until run-off at a concentration of 1 × 10^6^ Pn pycnidiospores ml^-1^ or 3000 Ptr spores ml^−1^ in 0.25% gelatin. The inoculated plants were kept in > 80% relative humidity for 24 h (Ptr inoculated plants) and 48 h (Pn inoculated plants) under a 12-h photoperiod and scored at 7 days post inoculation. A score of 1 to 9 was used to assess SNB disease response where 1 indicates no disease symptoms and a score of 9 signifies a fully necrotised plant as described by Phan et al. (2018). For TS disease phenotypes, a 1 to 5 scale was applied, in which 1 represents high resistance, and 5 represents high susceptibility (Lamari and Bernier 1989).

Field adult infection assay was conducted for SNB only. The DH105 x 56 lines were planted in hillplots of 8–10 seeds in each hill/line at Curtin University field trial site, Perth. The field trial was arranged with spacing of 30 cm between hills in all directions and bordered by two rows of a susceptible variety ‘Halberd’. The same pycnidiospore inoculum of Pn as described above was sprayed on 10-week-old plants until run-off. Fourteen days after the inoculation, disease severity was scored for flag leaves-1 (Flag-1), flag leaves-2 (Flag-2) and flag leaves-3 (Flag-3) as percentages of necrotic and/or chlorosis leaf-area (Eyal 1987). Within each hill plot, three individual plants were chosen randomly for disease scoring. This infection assay was conducted over winter growing season that is conducive for SNB disease development.

### Effectors and crude culture filtrates plant bioassays

To reduce genetic complexity in quantitative trait analysis, the double haploid DH105 x 56 lines that contain *Snn3* were excluded from the analysis. The selection was achieved by screening the 531 lines for sensitivity to SnTox3, using purified heterologous expressed protein. SnTox3 protein was produced in *Pichia pastoris* using pGAPzA expression system (Thermo Fisher Scientific, MA USA) as described by Phan et al. (2021). For mapping SnTox267 sensitivity loci, purified protein was expressed in *E. coli* SHuffle^®^ strain as described by Zhang et al. (2017) with some modifications. Briefly, Sn*Tox267* gene (GenBank accession QRD01761Q), which belongs to isoform group 2 (Supplementary Figure S1) (Richards et al. 2022) was amplified from the gDNA of SN15 isolate (Syme et al. 2013) using primer pair *CATATG*GCACCAGCCCCTGACAGC and *CTCGAG*GATACCGTGCCATGAGAGACGAG, and cloned into the expression vector pET21a(+) (Novagen). PCR amplification was performed in a 50 µl reaction containing 1 × Phusion HF master mix (NEB), 0.2 µM of each primer and 2 µl of 50ng/µl gDNA as template, and the annealing temperature used was 65°C. Protein was purified using HisTrap^™^ HF, 5ml (Cytavia), coupled to an AKTA purifier (GE Healthcare), and eluted with elution buffer (50 mm HEPES (*N*−2-hydroxyethylpiperazine-*N*’-2-ethanesulfonic acid) buffer pH 8.0, 300 mM NaCl and 100 mM imidazole). Purified protein was dialysed in 20 mM sodium phosphate buffer pH 7.0 and stored at −80°C until required for plant infiltrations.

Culture filtrate of Pn and Ptr isolates were produced in Fries 3 media as previously described (Liu et al. 2004; Moffat et al. 2014). The filtrate was harvested by filtration through sterile gauze, MiraCloth (Merck Millipore, Billerica, MA, USA), and passed through a 0.2-μm syringe filter unit (Pall Life Sciences, Port Washington, NY, USA). The expressed protein and culture filtrates were infiltrated into the first leaf of 2-week-old wheat seedlings using 1-mL needleless syringe. Plants were grown in vermiculite as described in the methodological section ‘Plant infection assays’ and maintained in a controlled growth chamber at 22 ± 1 °C with a 12-h photoperiod. The infiltrated leaves were visually evaluated for symptoms and scored on a scale of 0 to 4 where 0 indicates insensitivity (no reaction); 1, slight chlorosis; 2, extensive chlorosis; 3, extensive chlorosis with some necrosis; and 4, extensive necrosis. Bacterial cell extract derived from an *E.coli* strain harbouring pET21 vector was used as a negative control for infiltration assays. All infiltrations were carried out in biological triplicates.

### Genotyping, genetic linkage and QTL mapping

Genotypic data was generated from wheat DArT platform (Diversity Arrays Technology Pty Ltd, Australia; http://www.diversityarrays.com/) for the DH105 x 56 population. These markers were initially filtered with a maximum threshold of 95% reproducibility, 90% call rate for markers, and 20% missing values over samples. A final genetic linkage map was built using MultiPoint v. 3.2 software (MultiQTL Ltd, Institute of Evolution, Haifa University, Israel). Kosambi mapping function was used to convert the recombination frequencies into genetic distance (cM). Segregation distortion of the markers was checked against the expected Mendelian segregation ratio for co-dominant inheritance in a double haploid population using Chi-squared tests. This function is a built-in function in the MultiPoint program. One representative marker for each group of co-segregating markers was kept. Single markers showing significant segregation distortion were removed to avoid bias and false linkages. Groups of linked markers that were similarly distorted were accepted for linkage mapping and QTL analysis. Phenotypic data included effector-induced infiltration responses, disease responses at seedling stage under controlled conditions and percentage of green area of adult plants in the field trial. Genotype data were used for QTL analysis with MultiQTL software, version 2.5 (MultiQTL Ltd, Institute of Evolution, Haifa University, Israel). Significance levels for each QTL detected were determined with 1000 permutations, and the standard deviation for each QTL determined by 1000 Bootstraps. QTL with a LOD score ≥ 3.0 were declared as significant. In cases where multiple QTL hit the same location, QTL may be announced at LOD ≥ 2.5. DArT seq or SNP marker sequences flanking or located within QTL intervals were used to blast against wheat ‘Chinese Spring’ reference genome assembly (Consortium et al. 2018)(IWGSC RefSeq v2.0) from the Plant Ensemble website (http://plants.ensembl.org/Triticum_aestivum/Info/Index). Information obtained were used to anchor the DArTseq or SNP markers to the wheat ‘Chinese Spring’ reference genome assembly to allow comparison between studies.

### Allele stacking analysis

To determine the effect of different combinations of alleles associated with TS and SNB resistance found in the double haploid mapping population, the progenies were assigned to groups based on the number of resistance-associated alleles/QTL detected following disease assessments with Pn and Ptr isolates. These groupings were then used to study the effect of QTL stacking on TS and SNB disease severity. Statistical analysis of differences between the means of these groups (ANOVA) were carried out using R v4.5.2 (R Core Team 2021) and ranking among groups was done using R packages agricolae v1.4.0 (Mendiburu and Yaseen 2020). These were to assess whether wheat lines carrying different combinations of TS/SNB-related QTL would be more resistant or susceptible to the diseases and identify effective breeding targets. Significant differences between the groups were declared at *p* < 0.01.

### Fluorescence microscopic observations

Microscopic examinations of the infection processes by Pn and Ptr were conducted by setting up plant infection assays on wheat lines ‘56:ZWB11’ and ‘105:ZIF14’. A susceptible cultivar ‘Halberd’ was included in this experiment for comparison purposes. The inoculation preparation was as described in ‘Plant infection assay’ using GFP strains of Ptr and Pn derived from previous studies (See and Moffat 2023; Solomon et al. 2006). To test if needle-pricked wound facilitates spore penetration and disease development, a section of 2-week-old leaf was randomly pricked with a sterile needle before inoculation. A mock inoculation on the needle-pricked leaves was also included as a control. For fluorescence microscopy, the leaf segments were observed using a Nikon A1 laser scanning confocal microscopy and an Andor Dragonfly spinning disk confocal microscopy. To visualise the fluorescence expression of the GFP isolates, an optical filter with excitation at 488 nm and emission at 525/50 nm was used in Nikon A1, while Andor Dragonfly was equipped with laser excitation at 488 nm and emission filter at 521/38 nm. Plant chlorophyll fluorescence was detected using optical filter with excitation at 640 nm and emission at 700/755 nm while excitation in Andor Dragonfly was 637 nm and emission filter at 685/47 nm. Images were captured using NIS-Elements Software (Nikon, Japan) for Nikon A1 confocal system and Fusion 2.4 for Andor Dragonfly spinning disk confocal system. All images were further processed using Imaris imaging software ver 11.0 (Oxford Instruments).

## Results

### Identification of SNB and TS resistance sources

In our search for wheat lines that exhibit resistance to both SNB and TS diseases, we selected five well-performing disease resistant lines to wheat leaf blotches from the CAIGE national disease field evaluation program (https://www.caigeproject.org.au/) and conducted an initial disease assessment using 12 genetically diverse Pn isolates (Phan et al. 2020). Analysis of variance for the overall mean showed significant differences between the wheat lines (Fig. 1A) with ‘56:ZWB11’ displaying the highest resistance to SNB, while 104:ZIF14, 105:ZIF14 and 12:ZIZ11 were comparable with the Australian wheat cultivars and ‘Chinese Spring’. Next, ‘56:ZWB11’ and ‘105:ZIF14’ were then selected for TS disease assessment under controlled growth condition, against two Ptr isolates (race1 and race 2) (See et al. 2023). Both ‘56:ZWB11’ and ‘105:ZIF14’ had the lowest overall disease mean scores among the panel of Australian wheat cultivars tested, except wheat cultivar ‘Hydra’, which also displayed effective resistance to TS (Fig.1B). Effector profiling of ‘105:ZIF14’ to the known Pn and Ptr effectors indicated that this line was insensitive to ToxA, SnTox1, SnTox3, SnTox5 and ToxB but exhibited strong sensitivity to SnTox267, whereas ‘56:ZWB11’ was found to be sensitive to SnTox3 but displayed a milder sensitivity to SnTox267 (Supplementary Table S4). These two wheat lines were then used as parental lines to develop a double haploid wheat population DH105 x 56, which segregates for SnTox267 and SnTox3. To reduce the background noise caused by SnTox3-*Snn3* interaction, progenies that are sensitive to SnTox3 were removed from the population set via effector sensitivity screening. A final set of 241 double haploid lines were then used for genotyping and phenotyping.

**Fig. 1.**
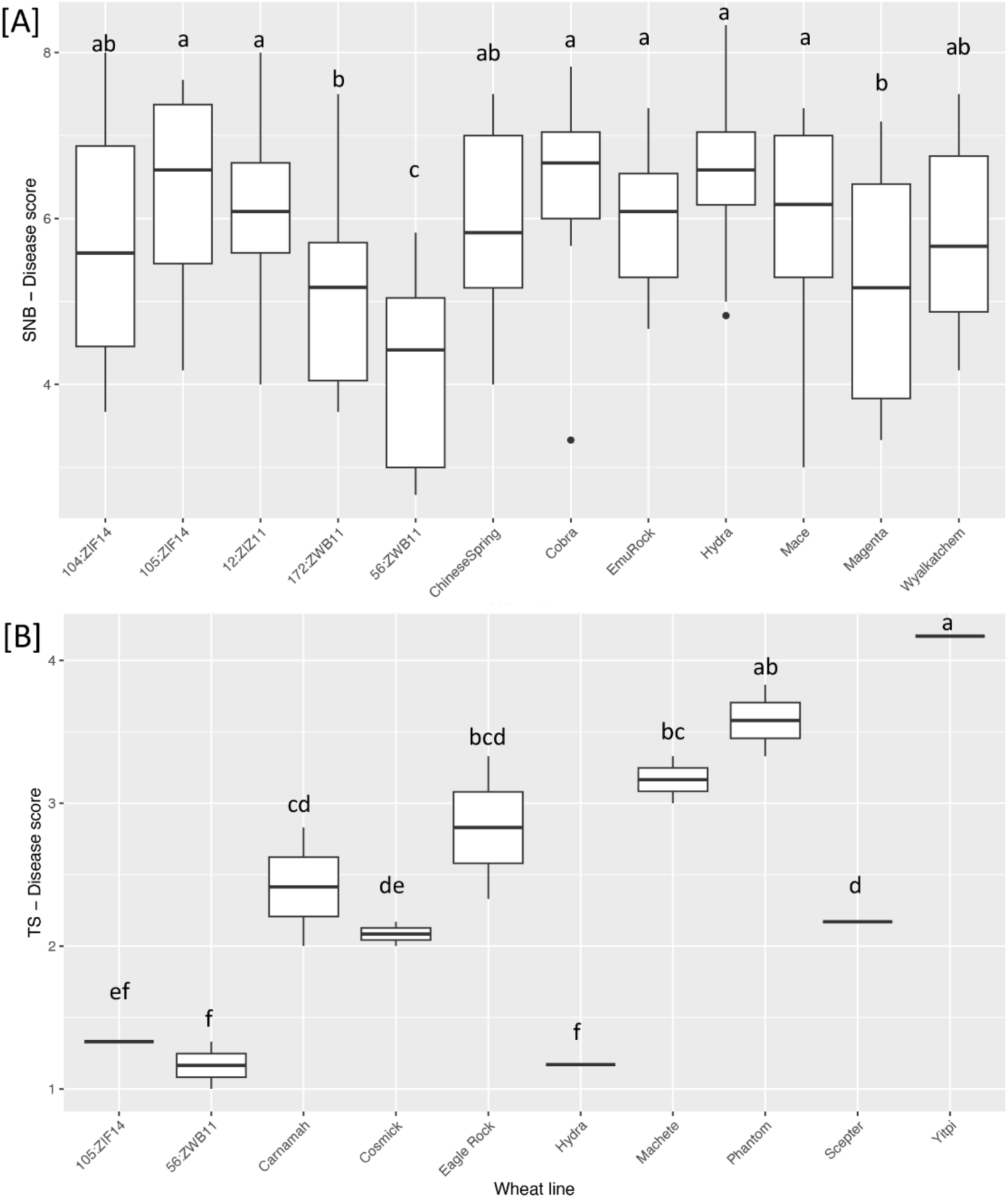
Septoria nodorum blotch (SNB) and tan spot (TS) disease responses for the CAIGE wheat lines in comparison to Australian wheat cultivars. Wheat cultivar ‘Chinese Spring’ was included in the SNB disease assessment. Overall mean disease response for each cultivar by a subset of (A) 12 *Parastagonospora nodorum* (Pn) isolates and (B) two *Pyrenophora tritici-repentis* (Ptr) isolates. Different letters above bars represent significant differences between treatments at a significance level of *p* < 0.05

### Genotyping and genetic linkage mapping

Wheat DArTseq platform was used to generate two large datasets of 12,225 DArTseq and 8,972 SNP markers for the DH105 x 56 population. Of these, 3,403 markers were successfully used to create 21 genetic-linkage maps representing 21 wheat chromosomes for the DH105 x 56 population (Table 1). The markers are evenly spaced across the whole genome with an average of 162.05 markers, 125.10 loci per chromosome and 3.49 cM interval (Table 1). The average of 416.19 cM length per linkage group (Table 1) indicates the presence of high level of recombination frequencies in the population.

**Table 1.** Genetic linkage map of DH105 x 56 double haploid wheat mapping population built with 3,403 DArTseq and SNP markers using MultiPoint software

| Linkage Group | Chromosome | Length (cM) | Number of loci | Number of makers | Density (cM) |
| --- | --- | --- | --- | --- | --- |
| LG1 | 1A | 424.31 | 143 | 173 | 2.97 |
| LG2 | 1B | 546.01 | 165 | 207 | 3.31 |
| LG3 | 1D | 356.83 | 80 | 104 | 4.46 |
| LG4 | 2A | 421.64 | 131 | 171 | 3.22 |
| LG5 | 2B | 575.03 | 175 | 233 | 3.29 |
| LG6 | 2D | 367.67 | 95 | 114 | 3.87 |
| LG7 | 3A | 454.90 | 165 | 222 | 2.76 |
| LG8 | 3B | 638.60 | 196 | 264 | 3.26 |
| LG9 | 3D | 413.65 | 109 | 144 | 3.79 |
| LG10 | 4A | 290.61 | 71 | 83 | 4.09 |
| LG11 | 4B | 240.40 | 94 | 114 | 2.56 |
| LG12 | 4D | 256.28 | 53 | 60 | 4.84 |
| LG13 | 5A | 417.69 | 138 | 181 | 3.03 |
| LG14 | 5B | 236.62 | 78 | 115 | 3.03 |
| LG15 | 5D | 309.50 | 62 | 75 | 4.99 |
| LG16 | 6A | 432.08 | 181 | 228 | 2.39 |
| LG17 | 6B | 357.23 | 119 | 156 | 3.00 |
| LG18 | 6D | 374.31 | 94 | 144 | 3.98 |
| LG19 | 7A | 640.49 | 213 | 273 | 3.01 |
| LG20 | 7B | 504.53 | 144 | 193 | 3.50 |
| LG21 | 7D | 481.67 | 121 | 149 | 3.98 |
| Total/Average | 21 | 416.19 | 125.10 | 162.05 | 3.49 |

### QTL mapping

Phenotype dataset of 12 traits were obtained by challenging the DH105 x 56 population to different treatments. These included host sensitivity responses to purified SnTox267 effector and crude culture filtrates (CF) from Pn and Ptr isolates, and plant infection assays at seedling stage using different SNB and TS isolates that represent the Australian pathotypes and field SNB disease assessment at adult stage. Histograms show a continuous distribution for both diseases and culture filtrates, an indication of multi-QTL traits (Supplementary Figure S2). QTL analysis was carried out using the phenotype dataset and the constructed genetic linkage map (Table 1). The analysis identified a total of 19 QTL across 11 chromosomes, and 68.42% of the QTL detected were associated with more than one trait (Table 2 and Fig. 1).

**Table 2.**
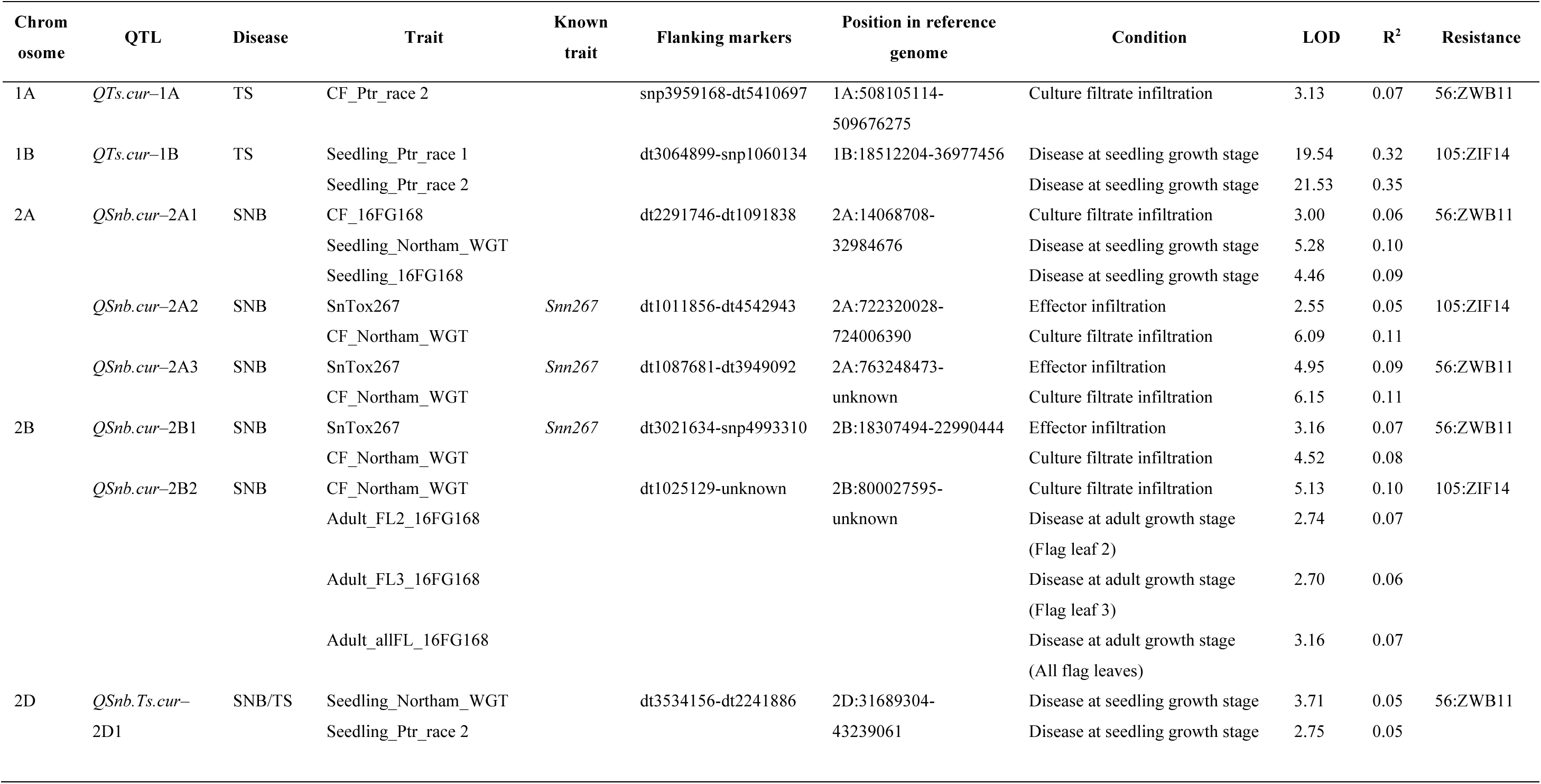
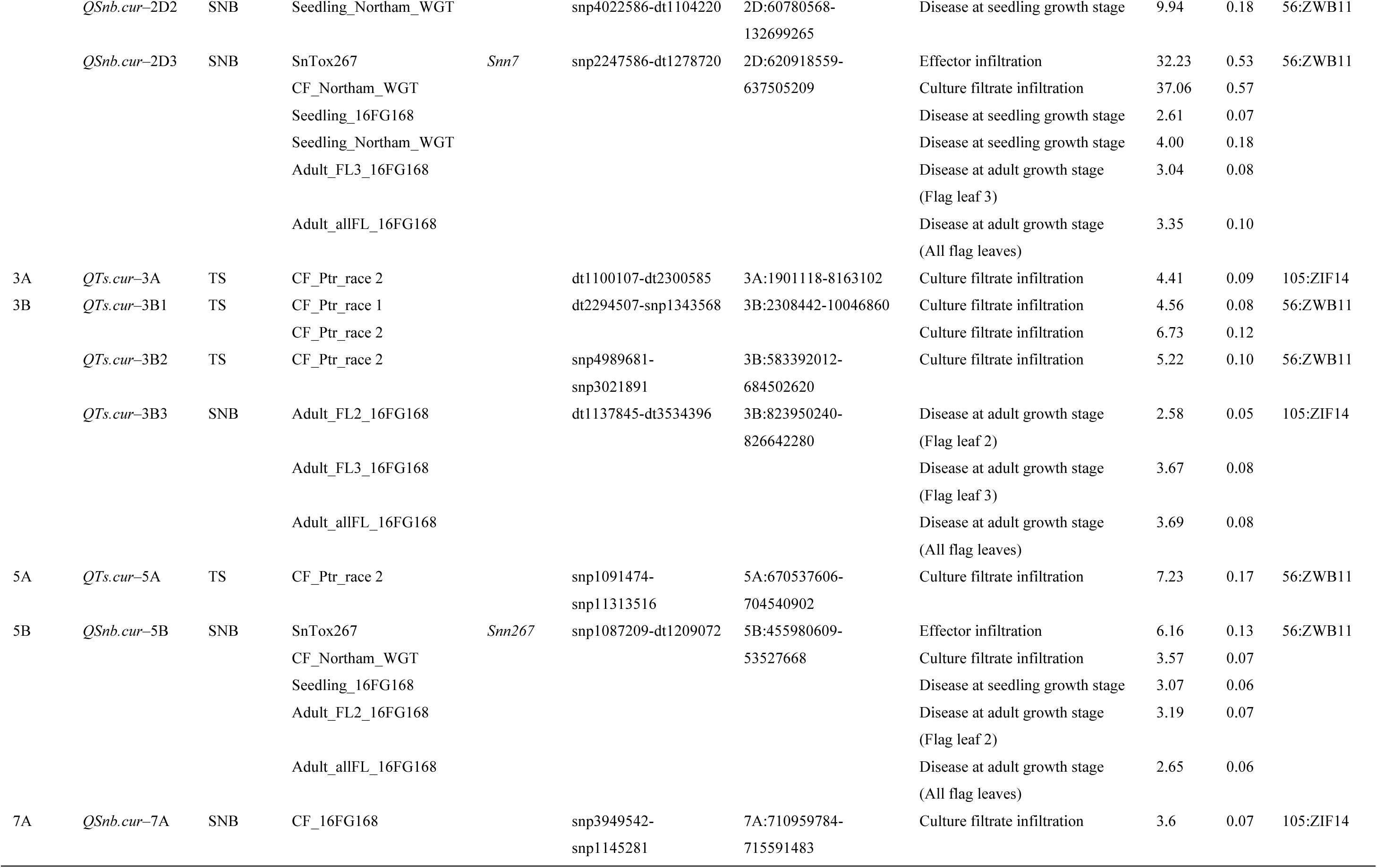
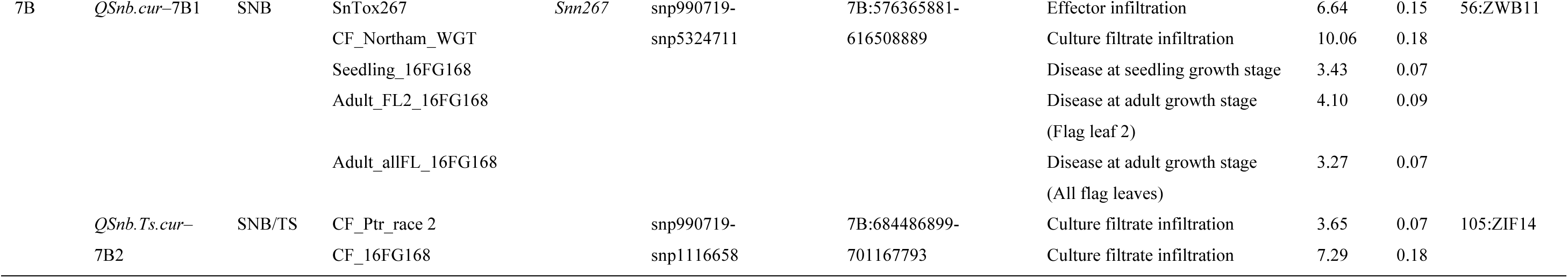
Summary of QTL detected in the DH105 x 56 population across 10 experiments including tan spot and septoria nodorum blotch disease assessments at seedling growth stage using two *Parastagonospora nodorum* and *Pyrenophora tritici-repentis* isolates of different pathogenicity profiles/races (Jones et al., 2024 and See et al., 2023), field disease assessment at adult growth stage (for SNB only), and sensitivity assays using culture filtrates from the respective isolates and purified effector SnTox267

### QTL for SNB

Six QTL located on chromosomes 2A (2A2, 2A3), 2B (2B1), 2D (2D3), 5B, and 7B (7B1) were detected for host response to purified SnTox267 effector. All six were found to be associated with host induced activity by culture filtrates of Pn isolate Northam_WGT and of those, three QTL (*QSnb.cur*–2D3, *QSnb.cur*–5B and *QSnb.cur*–7B1) were associated with SNB disease at both seedling and adult stages, explained by phenotypic variation ranging from 6% to 18% across the different experiments (Table 2, Fig. 2). For the culture filtrate of Pn isolate 16FG168 (CF_16FG168), two significant QTL were detected residing in chromosomes 2A (2A1) and 7B (7B2) (Table 2, Fig.2), where the QTL on 2A *QSnb.cur*–2A1 also co-localized with seedling disease response. No known SNB effectors was found mapped to these locations. For SNB disease, besides the three all-stage disease QTL (*QSnb.fcu*–2D3, *QSnb.cur*–5B and *QSnb.cur*–7B1) that correspond to SnTox267 sensitivity, additional two QTL (*QSnb.cur*–2B2 and *QTs.cur*–3B3) were associated with adult plant response, and three QTL (*QSnb.cur*–2A1, *QSnb.Ts.cur*–2D1 and *QSnb.cur*–2D2) with seedling response. All these stage-specific QTLs were minor effect QTL except *QSnb.cur*–2D2, which was associated with seedling disease response upon Pn Northam_WGT inoculation, explained by 18% phenotypic variation (Fig.2, Table 2). Singleton QTL were identified on chromosomes 2D (*QSnb.fcu*–2D2) and 7A (*QSnb.cur*–7A) that correspond to SNB disease at seedling stage detected by Pn isolate Northam_WGT and infiltration of culture filtrate derived from Pn isolate 16FG168, respectively (Table Fig. 2).

**Fig. 2.**
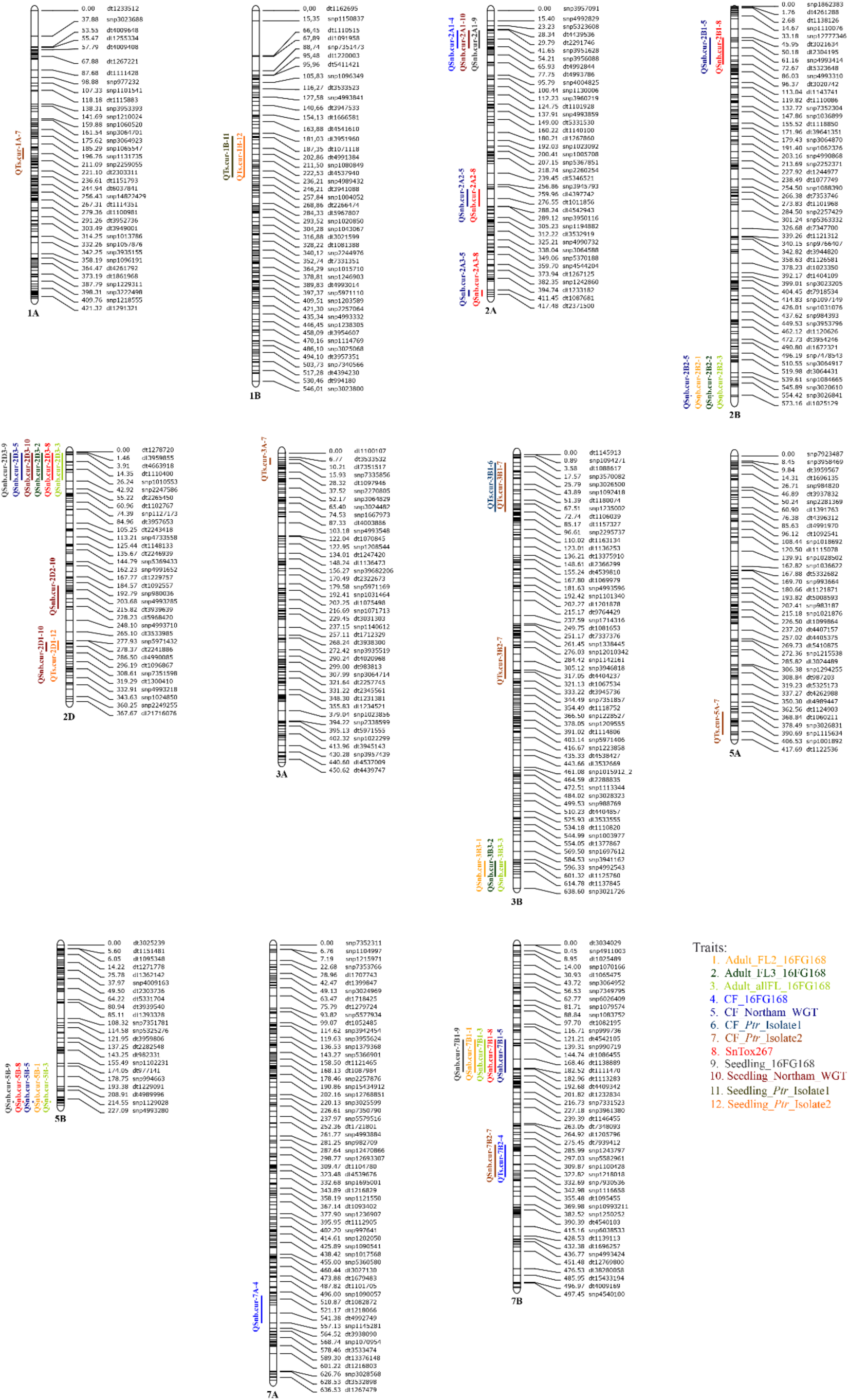
Map positions of quantitative trait loci (QTL) associated with septoria nodorum blotch and tan spot diseases, culture filtrates and SnTox267 effector sensitivity loci present on the genetic linkage map of the DH105 x 56 double haploid wheat population. Genetic maps of the chromosomes with trait markers highlighted on the left, and the genetic markers and the centimorgan (cM) distances between loci shown on the right. There are total of 19 QTL for 12 traits identified on 11 wheat chromosomes. The traits are colour coded as depicted in the figure legend. ‘CF’ and ‘FL’ denote culture filtrate and flag leaf, respectively

### QTL for Tan spot

Of the total 19 QTL detected in this study, two (*QSnb.Ts.fcu*–2D1 and *QSnb.Ts.fcu*–7B2) were common between TS and SNB-related traits. These QTL were associated with seedling resistance to Pn Northam WGT and Ptr race 2 inoculations, and host sensitivity responses to culture filtrates of Pn 16FG168 and Ptr race 2, respectively (Table 2). QTL profiles for the culture filtrate of the two Australian Ptr races were distinct, with six QTL (*QTs.cur*–1A, *QTs.cur*–3A, *QTs.cur*–3B1, *QTs.cur*–3B2, *QTs.cur*–5A and *QSnb.cur*–7B2) found to correspond to CF_Ptr_race 2, while only one of them (*QTs.cur*–3B1) was associated with CF_Ptr_race 1. Neither QTL was linked to known TS effectors. Intriguingly, TS seedling disease resistance was primarily controlled by a single major QTL on chromosome 1B (*QTs.cur*–1B), which came from the parent ‘105:ZIF14’ (Table 2, Fig. 2). Marker sequences surrounding this 1B QTL interval were used as queries in BLASTn searches of ‘Chinese Spring’ reference genome at Plant Ensemble (https://plants.ensembl.org/index.html) to determine if this QTL is linked to the 1BS TS resistance QTL reported in other studies (Dinglasan et al. 2019; Faris and Friesen 2005; Liu et al. 2020; Shankar et al. 2017; Taylor et al. 2023). Most of the closely linked markers did not have good matches to ‘Chinese Spring’, which suggests that ‘Chinese Spring’ does not carry this resistance allele; however, close-by markers have hits at the distal end of 1B chromosome short arm (Supplementary Table S5). This comparative analysis on chromosome 1B suggests that *QTs.cur*–1B is in the same region as previously identified 1BS TS resistance QTL.

### QTL effects

To assist breeders in making breeding decisions, we examined the effects of the identified single and stacked QTL in both disease outcomes. For a single-gene trait (Seedling_Ptr_race 1), we can observe a significant reduction in TS disease load with the presence of the resistance allele *QTs.cur*–1B from ‘105:ZIF14’ (Fig. 3A). For multiple-gene traits, higher resistance can be achieved by stacking higher numbers of genes/QTL, however there was a limit to the effectiveness. While additional QTL significantly improved seedling-stage disease response to TS (Fig. 3B), the cumulative effect was much more complex in SNB. Maximum effect of stacking multiple genes/QTL was observed with a combination of three or four genes/QTL for SNB seedling stage (Fig. 3C), however at seedling stage, significant effect was only detected for isolate Northam_WGT, not for isolate 16FG168 (data not shown). Meanwhile, four gene/QTL combination was the best for SNB resistance in field condition at adult stage (Fig. 3D).

**Fig. 3.**
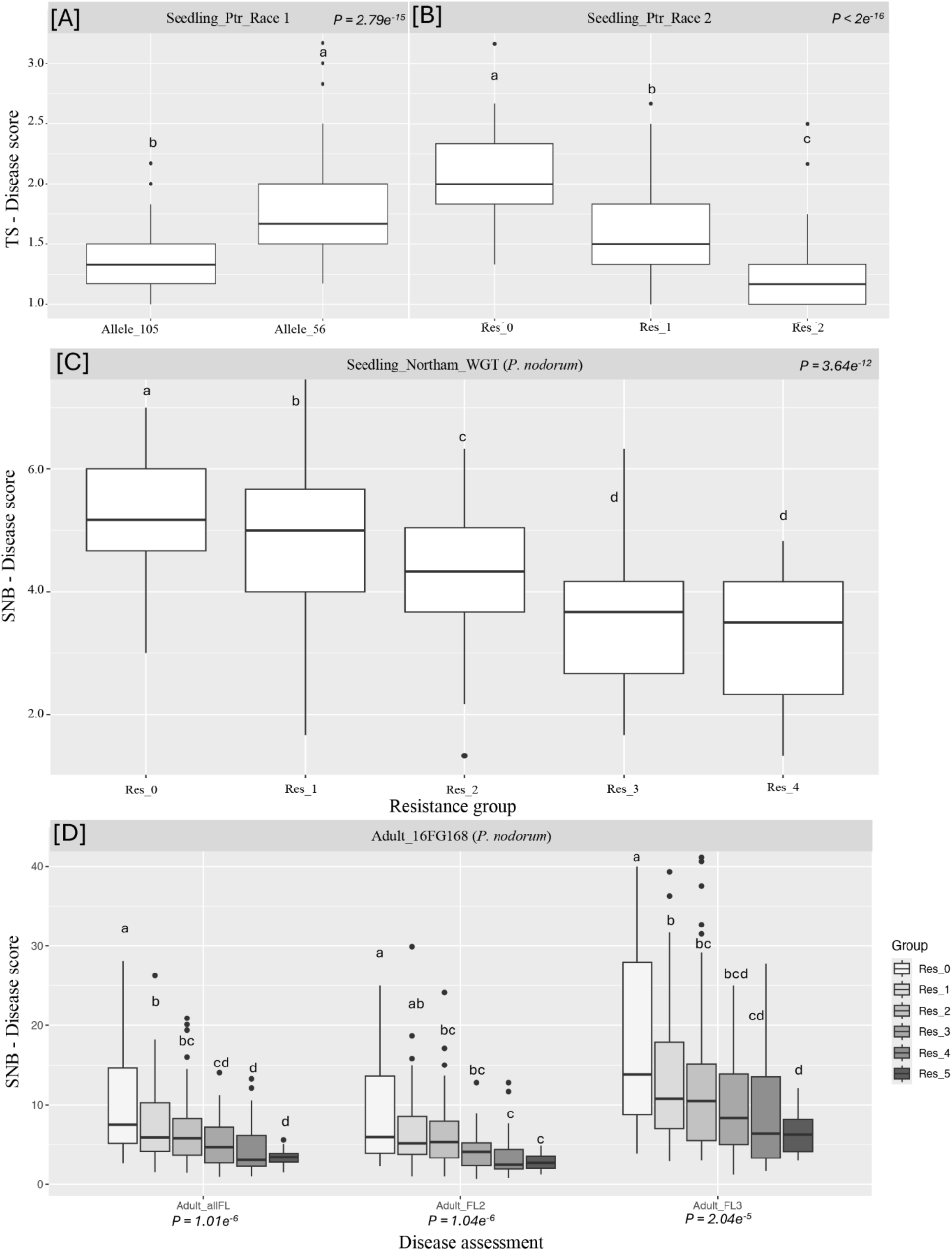
Effect of QTL stacking. Boxplots showing disease scores of progeny lines from DH105 x 56 population grouped based on resistance QTL numbers calculated across *Pyrenophora tritici-repentis* infections at seedling growth stage (A, B), *Parastagonosprora nodorum* infections at seedling (C) and adult growth stages (D). Res_0, Res_2, Res_3, Res_4 and Res_5 represent progeny lines carrying 0 to 5 QTL, respectively. Different letters indicate significant differences (ANOVA, *p* < 0.05)

### Microscopy visualisation of Pn and Ptr infection process

To further understand the infection process of Pn and Ptr on resistant genotypes, confocal microscopy was used to visualise SNB and TS disease developments on ‘56:ZWB11’ and ‘105:ZIF14’ at 24 h, 48 h and 96 h post-inoculation. A susceptible cultivar ‘Halberd’ was included as a control. To test the hypothesis that physical barriers may play a role as a resistance mechanism against Pn and Ptr, the wheat leaves were also wounded by pricking the leaf with a needle prior to inoculation, and visualised along with the unwounded leaves. At 24 h, Ptr conidia germinated on the leaf surface of the resistant and susceptible hosts to produce germ tube from basal cells (Fig. 4A, 4B and 4C). Germ tube extension however, varied between the hosts with longer germ tube extension observed on the susceptible ‘Halberd’ cultivar while Ptr conidia produced shorter germ tube on the resistant lines that even gave rise to club-shaped or appressorium-like structure on ‘56:ZWB11’ (Fig. 4A). Invasion of epidermal cells was evident only on the susceptible host (Fig. 4C). By 48 h, direct penetration of Ptr conidia was observed on all host leaf surfaces with intracellular mycelia growth more widespread on ‘Halberd’, followed by ‘105:ZIF14’ and ‘56:ZWB11’ (Fig. 4D, 4E and 4F). This disease symptom at micro-level on the resistant lines, however, was not evident to the naked eye in particular on ‘56:ZWB11’ where the leaves remained asymptomatic at 48 h (Fig. 5C). Leaf wounding did not facilitate Ptr conidium invasion on neither host although conidium hyphal extension and penetration were evident adjacent to the puncture (Fig. 4G, 4H and 4I). By 96 h, Halberd leaves had more extensive intracellular mycelia growth in comparison to ‘105:ZIF14’ and ‘56:ZWB11’ (Fig. 4J, 4K and 4L). This correlates with the classic tan spot disease lesions (and coalesced lesions) that were visible on the susceptible host (Fig. 5E), whereas both ‘105:ZIF14’ and ‘56:ZWB11’ produced small lesions with restricted mycelia growth even with the disruption of the leaf physical barrier (Fig. 4M and 4N). Intriguing, Halberd leaves that were not wounded displayed a more severe disease symptom in comparison to the treated leaves (Fig. 5E). This was also observed on ‘105:ZIF14’ but to a lesser extent of disease lesion development compared to ‘Halberd’. The treatment of needle-pricking on the leaf without conidium inoculation did not induce any symptoms for all time points investigated (Supplementary Figure S3B, 3D and 3F).

**Fig. 4.**
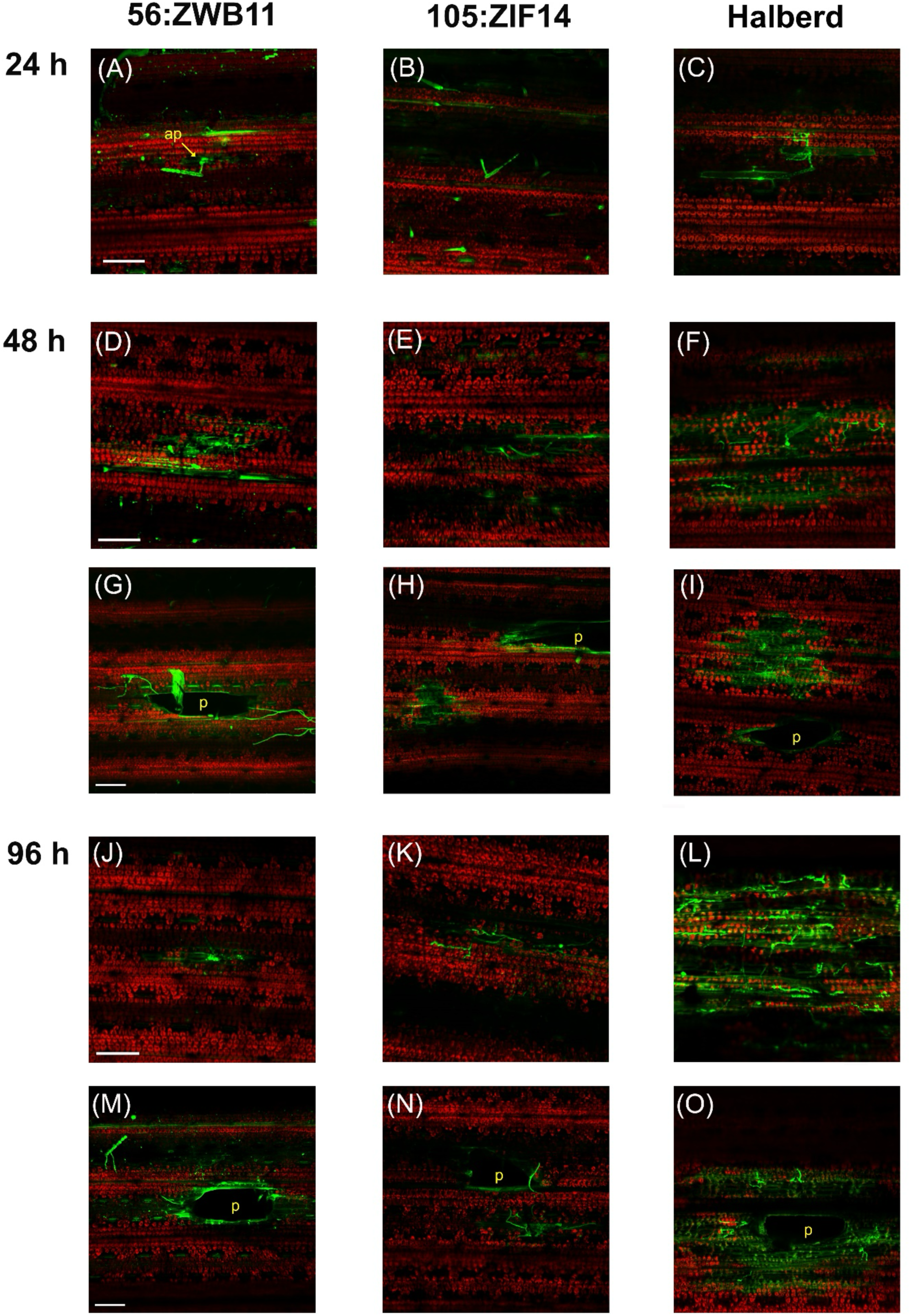
Confocal fluorescence microscopic examination of *Pyrenophora tritici-repentis* on resistant (‘56:ZWB11’ and ‘105:ZIF14’) and susceptible (‘Halberd’) cultivars of wheat. GFP strain conidia were inoculated on wheat leaves of ‘56:ZWB11’ (right panels), ‘105:ZIF14’ (middle panels) and ‘Halberd’ (left panels), and observed at 24 h (A - C), 48 h (D – I) and 96 h (J – O) post-inoculation. Fungal conidia and hyphae appear as bright fluorescence green and plant chlorophyll as bright fluorescence red. Damaged plant cells autofluorescence green under FITC fluorescence filters. ap, appressorium-like structure; p, puncture. Bar = 150 µm

**Fig. 5.**
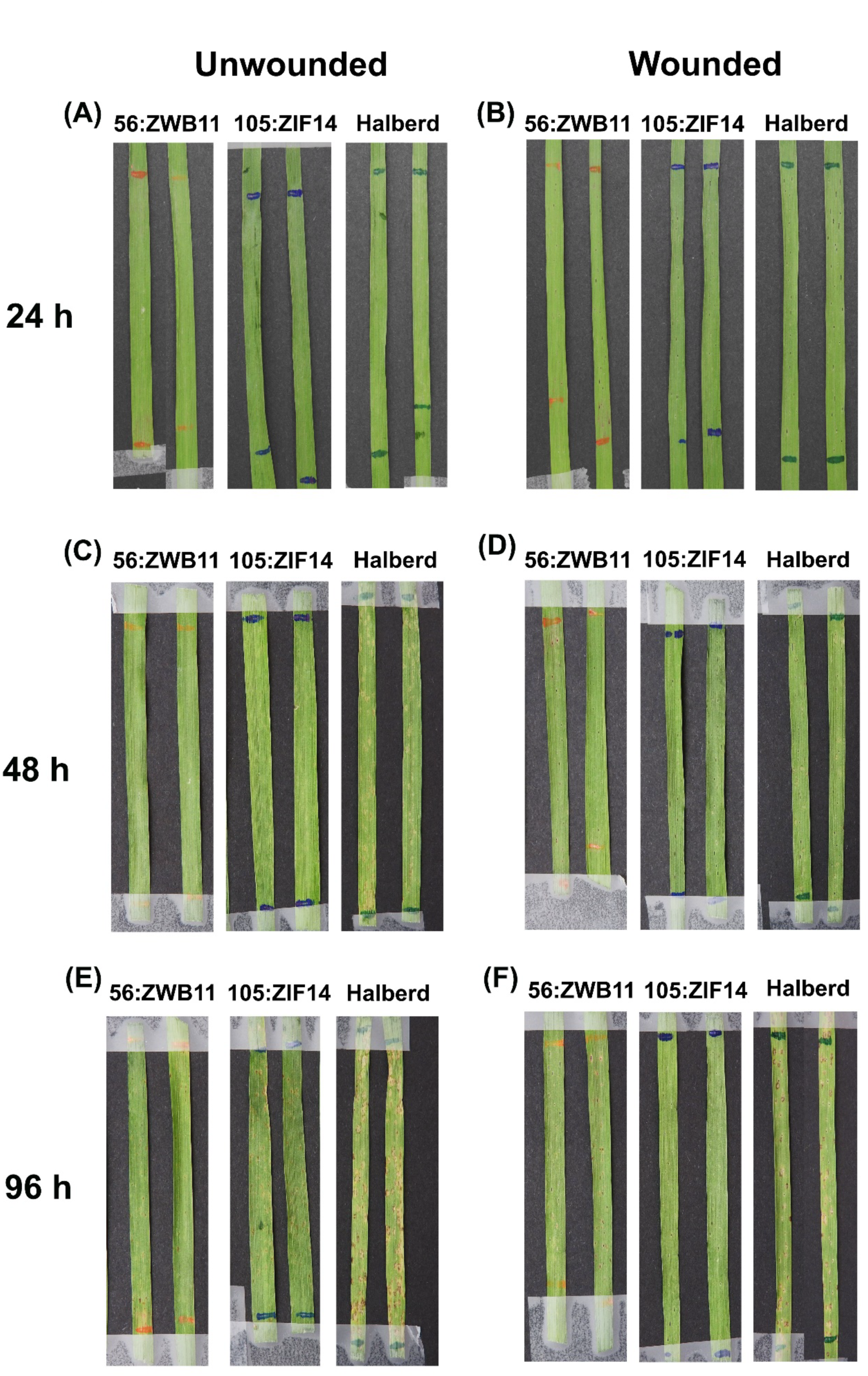
Tan spot disease symptoms development on wheat leaves upon a time course infection of *Pyrenophora tritici-repentis* GFP strain conidia. Unwounded and wounded (needle-prick) leaves were examined alongside at time-point 24 h, 48 h and 96 h. Resistant wheat lines - ‘56:ZWB11’ and ‘105:ZIF14’, susceptible control - ‘Halberd’

On Pn infected wheat leaves, germinated conidia were observed on all three wheat leaves at 24 h (Fig. 6A, 6B and 6C). The behaviour of the conidial extension did not differ between ‘56:ZWB11’, ‘105:ZIF14’ and ‘Halberd’. At 48 h, penetration attempts were generally associated with the resistance cultivars, including extension of germ tube through stomal subsidiary cells (Fig. 6G and 6H), while successful penetration into the cuticle cells were mainly observed in ‘Halberd’ with Pn hyphae extended across the anticlinal walls (Fig. 6I). This development, however, only appeared as microscopic symptoms on ‘Halberd’ wheat leaves as disease symptoms at this stage were barely visible to the naked eye (Fig. 7C). On the wounded leaves, hyphal colonisation was observed surrounding the puncture for all three wheat lines. Reduction of chloroplast cells peripheral to the wound (Fig. 6J, 6K and 6L) resulted in small chlorotic halo symptoms on ‘56:ZWB11’, ‘105:ZIF14’ and ‘Halberd’ (Fig. 7D). At 96 h, higher Pn hyphal density was observed surrounding the wound but with restricted lateral growth on ‘56:ZWB11’ (Fig. 6P). Despite Pn hyphae were seen to grow into the periclinal walls and the intercellular spaces of the mesophyll (Fig. 6P and 6Q), the disease symptoms on ‘56:ZWB11’ and ‘105:ZIF14’ leaves remained minuscule at 96 h (Fig. 7F). In contrast, an extensive network of Pn mycelia was observed in the leaf puncture of the susceptible line ‘Halberd’ (Fig. 6R) with lesions coalescing on the leaves (Fig. 7F). Similar disease symptoms were also observed on the unwounded ‘Halberd’ leaves (Fig. 7E). Under microscopic examination, widespread Pn mycelia growth accompanied by chloroplast degradation were evident (Fig. 6O). Colonisation of Pn on ‘Halberd’ leading to proliferation of hyphae through the stomata was also observed. On the resistant lines, there were no major tissue alterations or extensive mycelia growth, although Pn conidia continued to germinate, causing minimal cell damages (Fig. 6M and 6N; Fig. 7E).

**Fig. 6.**
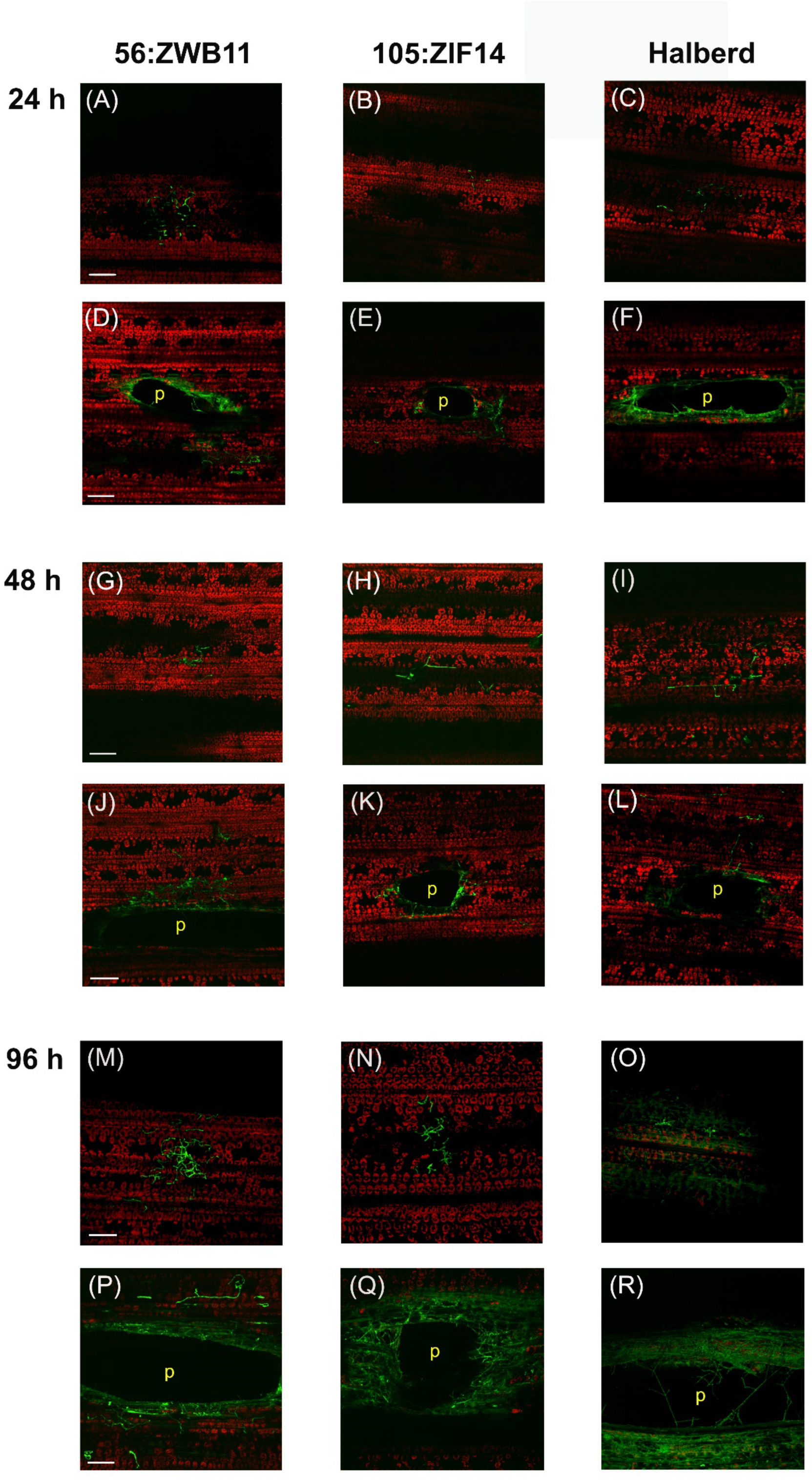
Confocal fluorescence microscopic examination of *Parastagonospora nodorum* on resistant (‘56:ZWB11’ and ‘105:ZIF14’) and susceptible (‘Halberd’) cultivars of wheat. GFP strain conidia were inoculated on wheat leaves of ‘56:ZWB11’ (right panels), ‘’105:ZIF14 (middle panels) and ‘Halberd’ (left panels), and observed at 24 h (A - F), 48 h (G – L) and 96 h (M – R) post-inoculation. Fungal conidia and hyphae appear as bright fluorescence green and plant chlorophyll as bright fluorescence red. Damaged plant cells autofluorescence as green under FITC fluorescence filters. ap, appressorium-like structure; p, puncture. Bar = 150 µm

**Fig. 7.**
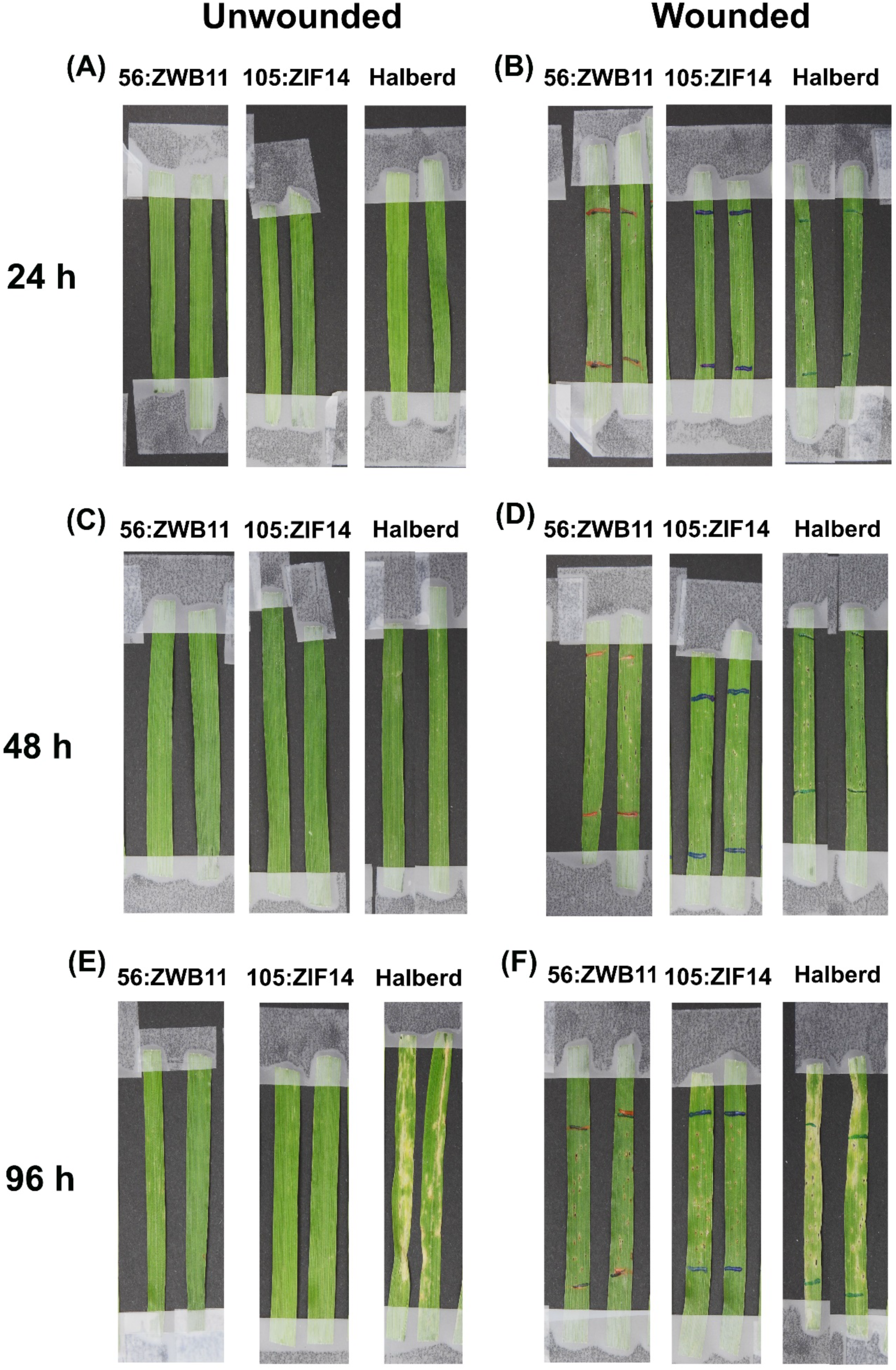
Septoria nodorum blotch disease symptoms development on wheat leaves upon a time course infection of *Parastagonospora nodorum* GFP strain conidia. Unwounded and wounded (needle-prick) leaves were examined alongside at time-point 24 h, 48 h and 96 h. Resistant wheat lines - ‘56:ZWB11’ and ‘105:ZIF14’, susceptible control - ‘Halberd’

## Discussion

Numerous studies have shown that Pn and Ptr produce multiple pathogen-secreted NEs that interact with host receptor-genes in an inverse gene-for-gene manner, a prominent concept in necrotrophic fungal pathology that has been extensively reviewed (Downie et al. 2021; Kariyawasam et al. 2023; Liu et al. 2017; Oliver et al. 2012; Peters Haugrud et al. 2022); while limited studies on novel resistance sources have also been reported (Faris et al. 2020; Phan et al. 2018). Here, we endeavour to identify and understand how resistance mechanisms function against these two important wheat diseases. Using representative pathotypes, this study revealed a complex profile for each disease, with most traits observed to be governed by multi gene interactions. Nineteen QTL across 11 chromosomes were identified, of which a high percentage (> 68%) were associated with two or more traits. SNB is a more complicated disease with almost twice as many QTL identified for SNB-related traits (11 SNB-specific QTL) in comparison to TS (6 TS-specific QTL). The genetic resistance to SNB and TS were distinct and mainly derived from separate sources. For TS, the major resistance *QTs.cur*–1B QTL, originated from ‘105:ZIF14’ accounted for more than 30% of phenotypic variance to both Ptr races present in Australia (See et al. 2023). A significant QTL on chromosome 1B *QTs.fcu*-1BS along with another QTL on chromosome 3B *QTs.fcu*-3BL were the first race-nonspecific resistance QTL reported for TS, identified in a population of recombinant inbred lines derived from spring wheats ‘Grandin’ and ‘BR34’ (Faris and Friesen 2005). From our comparative analysis, it is possible that the *QTs.cur*–1B that confers resistance to Ptr race 1 and race 2 is the same with the race-nonspecific *QTs.fcu*-1BS. Reports of QTL 1B that confer resistance to TS were not limited to disease assessment in controlled environments but also included Australian field evaluations (Shankar et al. 2017; Taylor et al. 2023), though it cannot be ascertained whether resistance were race 1 and/or race 2 specific under these conditions. A meta-QTL analysis of TS resistance by Liu et al. (2020) assigned 1B as one of the meta-QTL, and candidate genes underlying the physical interval included several NBS-LRR resistance genes and powdery mildew *Pm3*-like resistance gene. A strong resistance source is invaluable and can be utilised in Australian breeding programs to develop TS resistant cultivars. Moreover, QTL or genomic blocks as such that confers resistance to both necrotrophic and biotrophic pathogens are highly sought after by crop breeders.

Among the wheat diseases caused by necrotrophic fungal pathogens, SNB is regarded as a highly complex pathosystem. Pn utilises a suite of effectors to manipulate host immune responses, and this includes the recently discovered SnTox267 (Richards et al. 2022). While SnTox267 has previously been reported to interact with host sensitivity genes located on three different loci, one on chromosome 6A (*Snn6*-SnTox6) and two on chromosome 2D (*Snn2*-SnTox2 and *Snn7*-SnTox7) (Friesen et al. 2007; Gao et al. 2015; Shi et al. 2015), in this study, five new QTL (*QSnb.cur*-2A2, *QSnb.cur*-2A3, *QSnb.fcu*-2B1, *QSnb.cur*-5B and *QSnb.cur*-7B1) were found to be associated with SnTox267 sensitivity. Of the three previously identified QTL, only *QSnb.fcu*–2D3 was mapped to a known position in the DH105 x 56 population, and that location is *Snn7* (Shi et al. 2015). Interesting to note that *QSnb.cur*–5B and *QSnb.cur*–7B1 were associated with SNB disease at both seedling and adult stages, albeit of minor QTL effects. These additional QTL found associated with SnTox267 is in agreement with Richards et al. (2022) study, in which the author postulated there could be additional host targets besides *Snn2*, *Snn6* and *Snn7*. Our observations support the notion that SnTox267 has the propensity to interact with multiple host targets present in wheat germplasm; SnTox267 plays the key role in inducing SNB disease load in conditions without SnToxA/1/3. Nevertheless, despite the disease complexity, this study demonstrated that resistance to SNB can be improved by stacking up to four resistance alleles.

Discovery of NE is notoriously tricky due to their lack of conserved functional domains, hence evaluating host sensitivity to fungal culture filtrate is often conducted as preliminary assessments to confirm presence of novel NE and if the NE-host interaction poses any significance in disease development. Here, the culture filtrate of different Pn and Ptr isolates showed distinct QTL profiles. Of the two Pn isolates, Northam_WGT was shown to produce SnTox267 *in vitro* through the co-detection of similar QTL with the purified SnTox267, meanwhile, the genetic interactions induced by 16FG168 culture filtrate were unique to SnTox267 and Northam_WGT culture filtrate. Of the three unique QTL, *QSnb.cur*–2A1was associated with SNB disease at seedling stage, suggesting the presence of a novel NE expressed only in the culture filtrate of isolate 16FG168. On the contrary, although there were QTL identified associated with the culture filtrates of both Ptr isolates, none was co-localized to a disease trait on DH105 x 56 population. Nonetheless, our QTL analysis identified a common QTL induced by the culture filtrates of Ptr and Pn, thus suggesting *QSnb.Ts.fcu*–7B2 may correspond to an unknown NE found in both species; this proposition however remains to be investigated. Altogether these results highlight the need for continuous research in identifying suitable set of representative isolates in parallel with germplasm development for effective disease screening.

Thus far, our genetic analysis has demonstrated that the underlying genetic basis of resistance/susceptibility in SNB and TS are discrete. This observation was supported by the cytological examinations using fluorescence microscopy. For SNB, its resistance component seems to be associated with epidermal or leaf surface-related protection, which might include physical or chemical defences against Pn. In this study, we observed that pricking the leaves of these resistant cultivars facilitates hyphal colonisation, a stark contrast to the susceptible cultivar ‘Halberd’ where high infection was observed in both intact and physically compromised leaves. Numerous studies have shown that plant cell wall actively remodelled and restrengthened specifically at the sites of contact with pathogens. This active reinforcement of the cell wall was reported to occur as an early response to perception of various types of pathogens including fungi (Underwood 2012). The cell wall reinforcement achieved through deposition of appositions, referred to as papillae, was also found to correlate with resistance to fungal pathogens (Böhlenius et al. 2010; Bruce 2014; von Röpenack et al. 1998). In contrast to SNB, high level of TS resistance was apparent in the wounded leaves. This suggests a completely different resistance mechanism beyond the leaf physical barrier. The higher resistance displayed in wounded leaves when inoculated with Ptr may implicate a certain level of basal defence response or “resistance priming” resulting in the wounded plants to be less susceptible. When plant physical barriers are breached, they are known to induce a plethora of defence responses including activation of stress-signalling pathways (jasmonic acid (JA) and salicylic acid (SA) signalling pathways), oxidative burst, defence-related pathways and cell wall reinforcement (Böhlenius et al. 2010; Savatin et al. 2014; von Röpenack et al. 1998). It would be of interest in further work on investigating the compounding factors that are involved in such resistance.

Combining different innate resistance mechanisms from different sources would be invaluable approach in breeding resistance to SNB and TS. Further investigation into the molecular mechanisms of the identified distinct resistance from ‘56:BWZ11’ and ‘105:ZIF14’ would be beneficial for precision breeding and our understanding on how resistance operates against these necrotrophic fungal pathogens. Meanwhile, germplasm of 24 double haploid progenies with optimal recombination carrying four or more SNB resistance QTL and TS resistance *QTs.cur*–1B (Supplementary Table S6) developed in this study, could provide valuable resources for integration into commercial elite lines.

Removal of the ToxA sensitivity gene *Tsn1* in breeding programmes has proven to be an effective strategy for improving TS resistance in Australia (See et al. 2024); however there are still a myriad of novel NEs to be uncovered in both pathogens. Removal of susceptibility by the elimination of effector sensitivity loci coupled with the integration of the above resistance resources will offer a sustainable solution to manage these complex fungal wheat diseases.

## Acknowledgements

This work was supported by the Grains Research and Development Corporation (GRDC) and Curtin University (project code CUR00023). We would like to thank Dr. Manisha Shankar and Dr. Hossein Golzar (Department of Primary Industries and Regional Development, DPIRD) for provision of seeds, Michael Nesbit and Patrick Lim (Curtin Health Innovation Research Institute, CHIRI) for technical support in fluorescence microscopy, and CAIGE (CIMMYT Australia ICARDA Germplasm Evaluation) program for providing germplasms and field evaluation data.

## Data availability statements

The datasets generated for this study can be found as supplemental data.

## Author contribution statement

HP conceived the experiment and wrote initial draft of the manuscript. PTS edited and wrote the manuscript. HP, EF and PTS designed the experiments. EF, FK, KR, LL, PTS, CC and KM performed the experiments. HP analysed the data. All authors reviewed and approved the manuscript.

## Ethics approval

On behalf of all authors, the corresponding author states that these experiments comply with the ethical standards in Australia.

