## Supplementary Figure S1 to S3 for "Differential genetic resistance identified in *Parastagonospora nodorum* and *Pyrenophora tritici-repentis*-wheat pathosystems"

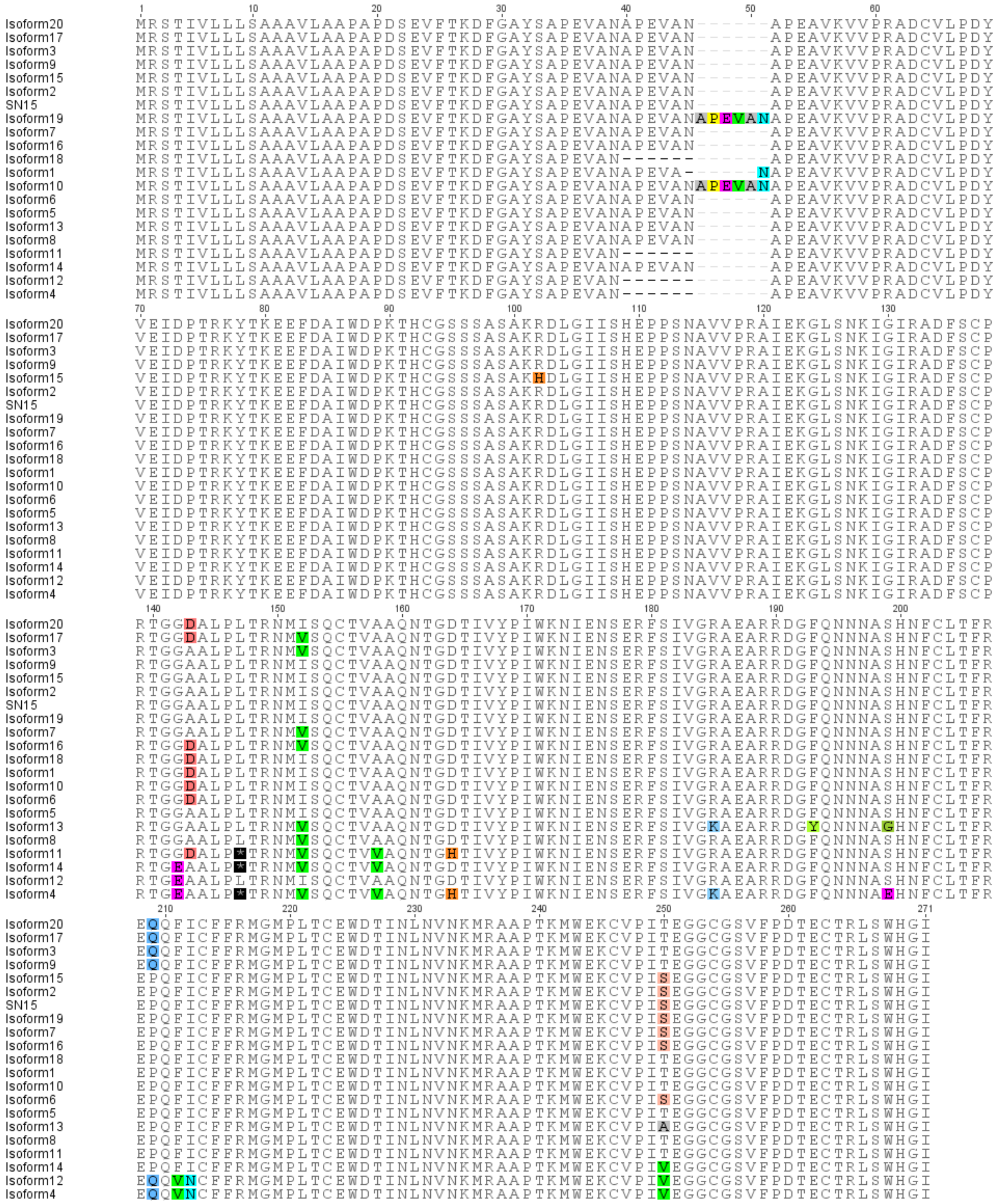

**Fig. S1** Amino acid alignment of SN15 SnTox267 with previously identified 20 protein isoforms (Richards et al., 2022). Coloured boxes represent non-conserved residue positions. A dash '-' represents a gap. The amino acids of the 20 SnTox267 isoforms were obtained from Richard et al. 2022.

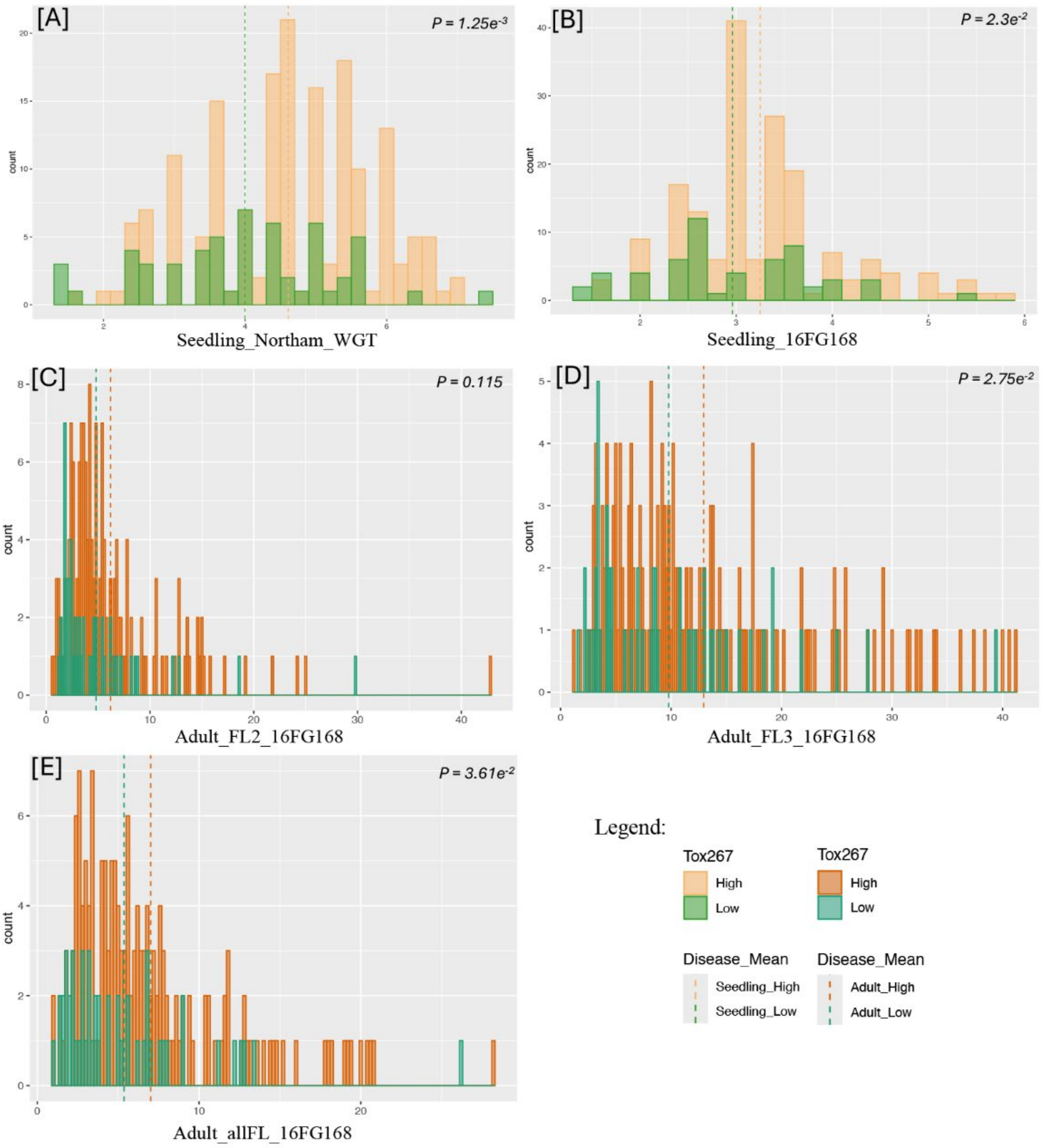


**Fig. S2** Histograms of septoria nodorum blotch diseases reactions of the DH105 x 56 double haploid recombinant population upon inoculation with *Parastagonospora nodorum* isolates Northam_WGT at seedling growth stage (A) and 16FG168 at seedling and adult growth stages (B – E). Sensitivity ratings of the DH105 x 56 lines to SnTox267 were categorized as either ‘high’ or ‘low’.

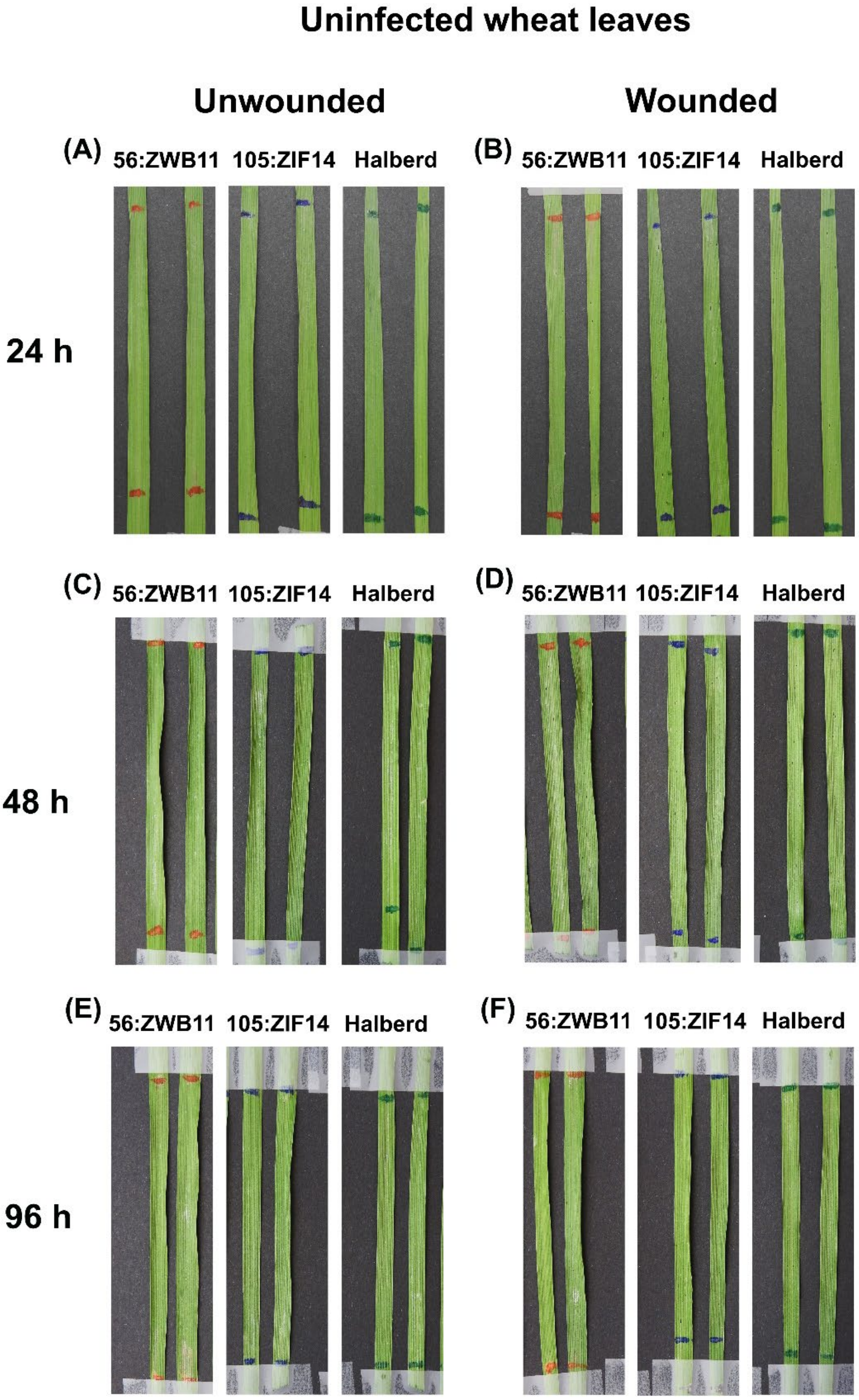
Uninfected wheat leaves
Unwounded
Wounded
(A)
56:ZWB11
105:ZIF14
Halberd
(B)
56:ZWB11
105:ZIF14
Halberd
24 h
(C)
56:ZWB11
105:ZIF14
Halberd
(D)
56:ZWB11
105:ZIF14
Halberd
48 h
(E)
56:ZWB11
105:ZIF14
Halberd
(F)
56:ZWB11
105:ZIF14
Halberd
96 h

**Fig. S3** Absence of disease symptom development on wheat leaves following a time course observation after spraying with 0.25% gelatine as control. Unwounded (A, C and E) and wounded (needle-prick) (B, D and F) leaves were examined alongside fungal inoculated leaves at time-point 24 h, 48 h and 96 h. Resistant wheat lines - '56:ZWB11' and '105:ZIF14', susceptible control - 'Halberd'
