## Supplementary Table S1 to S5 for "Differential genetic resistance identified in *Parastagonospora nodorum* and *Pyrenophora tritici-repentis*-wheat pathosystems"

**Table S1** Wheat germplasm from the CIMMYT (International Maize and Wheat Improvement Centre) Australia ICARDA (International Centre for Agriculture in the Dry Areas) Germplasm Evaluation (CAIGE) program

| Line | Source | Pedigree |
|---|---|---|
| 104:ZIF14 | ICARDA | SEKSAKA-6/IZAZ-8 |
| 105:ZIF14 | ICARDA | SEKSAKA-6/IZAZ-8 |
| 12:ZIZ11 | ICARDA | Not available |
| 56:ZWB11 | CIMMYT | Not available |
| 172:ZWB11 | CIMMYT | Not available |

**Table S2** Australian wheat cultivars used in this study

| Cultivar name | Year of release | Breeder |
|---|---|---|
| Carnamah | 1996 | Department of Agriculture and Food, Western Australia |
| Cobra | 2011 | LongReach Plant Breeders |
| Cosmick | 2014 | InterGrain |
| Eagle Rock | 2004 | Enterprise Grains Australia |
| Emu Rock | 2011 | InterGrain |
| Hydra | 2014 | InterGrain |
| Mace | 2008 | Australian Grain Technologies |
| Machete | 1985 | South Australia Research and Development Institute |
| Magenta | 2007 | InterGrain |
| Phantom | 2012 | LongReach Plant Breeders |
| Scepter | 2015 | Australian Grain Technologies |
| Wyalkatchem | 2001 | InterGrain |
| Yitpi | 1999 | South Australia Research and Development Institute |

**Table S3** The panel of 12 *Parastagonospora nodorum* isolates derived from Phan et al. (2020) used in this study to assess the five CAIGE wheat lines

| Isolate ID | Location | Year |
|---|---|---|
| 15FG114 | Southern Brook | 2014 |
| 15FG119 | Southern Brook | 2014 |
| 15FG49 | Muresk | 2015 |
| RAC2182_2 | Northam | 2015 |
| SN15 | Western Australia | 2001 |
| WAC13071 | Geraldton | 2005 |
| WAC13073 | Geraldton | 2005 |
| WAC13524 | Geraldton | 2011 |
| WAC13532 | Geraldton | 2011 |
| WAC13617 | Geraldton | 2012 |
| WAC8384 | Geraldton | 1990 |
| WAC8410 | Badgingarra | 1991 |

**Table S4** Effector sensitivity profiles of the two parental wheat lines ('105:ZIF14' and '56:ZWB11') and effector profiles of the *Parastagonospora nodorum* and *Pyrenophora tritici-repentis* fungal isolates used in this study

| Organism | Description | ToxA | SnTox1 | SnTox3 | SnTox267 | SnTox5 | ToxB |
|---|---|---|---|---|---|---|---|
| Wheat | '56:ZWB11' effector sensitivity score | 0.0 | 0.0 | 3.3 | 1.0 | 0.0 | 0.0 |
| | '105:ZIF14' effector sensitivity score | 0.0 | 0.0 | 0.0 | 4.0 | 0.0 | 0.0 |
| | '56:ZWB11' effector sensitivity rating | Insensitive | Insensitive | Sensitive | Mild sensitivity | Insensitive | Insensitive |
| | '105:ZIF14' effector sensitivity rating | Insensitive | Insensitive | Insensitive | Strong sensitivity | Insensitive | Insensitive |
| *Parastagonospora* | Northam_WGT | Presence | Presence | Presence | Presence | Presence | NA |
| *nodorum* | 16FG168 | Presence | Presence | Presence | Presence | Presence | NA |
| *Pyrenophora* | Ptr_race 1 | Presence | NA | NA | NA | NA | Absence |
| *tritici-repentis* | Ptr_race 2 | Presence | NA | NA | NA | NA | Absence |

NA – not applicable

**Table S5** Comparative mapping of 1BS QTL for resistance to tan spot

| Population | QTL name | Phys. interval (Mbp)[1] | Gen.map. interval (cM) | Ave. LOD/-log10(P) | Ave. %GV / Cumulative Marker Effect[2] | Trait | Source of allele that reduces trait value | Reference |
|---|---|---|---|---|---|---|---|---|
| DH105 x 56 | *QTs.cur*–1B | 18.5-37 | 185-223 | 21.53 | 0.35 | Race1 and race 2, seedling | 105:ZIF14 | This study |
| IGW2574/Annuello | 1B | NA | 13.5 – 14.8 | 2.2 | 9.00 | Tan spot severity (booting) | IGW2574 | Shankar et al. (2017) |
| BR34/Grandin | *QTs.fcu*-1BS | 3.6–6.3 | 13.9 ~ 30.1 | 5.47 | 14.00 | 86-124 (race 2), seedling | | Liu et al. (2020); Faris and Friesen (2005) |
| BR34/Grandin | *QTs.fcu*-1BS | 3.6–6.3 | 13.9 ~ 30.1 | 5.94 | 13.00 | DW5 (race 5), seedling | | Liu et al. (2020); Faris and Friesen (2005) |
| BR34/Grandin | *QTs.fcu*-1BS | 3.6–6.3 | 13.9 ~ 30.1 | 8.61 | 29.00 | OH99 (race 3), seedling | | Liu et al. (2020); Faris and Friesen (2005) |
| BR34/Grandin | *QTs.fcu*-1BS | 3.6–6.3 | 13.9 ~ 30.1 | 8.05 | 27.00 | Pti2 (race 1), seedling | | Liu et al. (2020); Faris and Friesen (2005) |
| 192 international wheat diversity panel | TQTL-1B.1 | 6.9–44.5 | 8.36 – 53.61 | 12.83 | 8.27 | Field evaluation (adult) | | Taylor et al. (2023) |
| 295 bread wheat collections | 1B | 32.46 | 51.29 | 3.17 | 0.991/-0.634 | Race 1, seedling | | Dinglasan et al. (2019) |

[1]Physical interval based on Chinese Spring reference genome from marker sequences

[2]Effect of the minor allele on the disease severity
