## Supplementary Table S6 for "Differential genetic resistance identified in *Parastagonospora nodorum* and *Pyrenophora tritici-repentis*-wheat pathosystems"

**Table S6:** QTL stacking identified 24 DH progeny resistant to both disease (SNB & TS)

| DH105x56 | QTs.cur–1B_105 | Group_SNB_Adult | Res_both_SNB_TS |
|---|---|---|---|
| H001 | Allele_56 | Res_3 | |
| H003 | Allele_56 | Res_2 | |
| H004 | Allele_56 | Res_4 | |
| H006 | Allele_56 | Res_3 | |
| H009 | Allele_105 | Res_3 | |
| H010 | Allele_56 | Res_2 | |
| H011 | Allele_56 | Res_3 | |
| H012 | Allele_56 | Res_1 | |
| H013 | Allele_56 | Res_0 | |
| H014 | Allele_105 | Res_0 | |
| H017 | Allele_105 | Res_4 | 1 |
| H018 | Allele_105 | NA | |
| H019 | Allele_105 | Res_2 | |
| H033 | Allele_105 | Res_2 | |
| H036 | Allele_56 | Res_0 | |
| H040 | Allele_105 | NA | |
| H043 | Allele_105 | Res_0 | |
| H047 | Allele_105 | Res_1 | |
| H052 | Allele_105 | Res_3 | |
| H054 | Allele_56 | Res_3 | |
| H056 | Allele_56 | Res_0 | |
| H058 | Allele_56 | Res_0 | |
| H061 | Allele_105 | Res_3 | |
| H063 | Allele_105 | Res_1 | |
| H064 | Allele_105 | Res_2 | |
| H065 | Allele_56 | Res_2 | |
| H067 | Allele_56 | Res_4 | |
| H072 | Allele_105 | Res_4 | 1 |
| H073 | Allele_56 | Res_3 | |
| H075 | Allele_105 | Res_4 | 1 |
| H078 | Allele_56 | Res_1 | |
| H081 | Allele_56 | Res_4 | |
| H082 | Allele_105 | Res_0 | |
| H087 | Allele_105 | Res_1 | |
| H088 | Allele_105 | Res_1 | |
| H089 | Allele_56 | Res_2 | |
| H091 | Allele_56 | Res_5 | |
| H093 | Allele_56 | Res_3 | |
| H095 | Allele_56 | Res_1 | |
| H096 | Allele_105 | Res_1 | |

| | | | |
|---|---|---|---|
| H098 | Allele_105 | Res_5 | 1 |
| H102 | Allele_105 | Res_0 | |
| H112 | Allele_56 | Res_3 | |
| H115 | Allele_56 | Res_1 | |
| H116 | Allele_56 | Res_1 | |
| H118 | Allele_105 | Res_0 | |
| H120 | Allele_56 | Res_3 | |
| H122 | Allele_56 | Res_1 | |
| H123 | Allele_56 | Res_5 | |
| H124 | Allele_56 | Res_3 | |
| H126 | Allele_56 | Res_3 | |
| H128 | Allele_56 | Res_3 | |
| H130 | Allele_105 | Res_2 | |
| H131 | Allele_56 | Res_1 | |
| H132 | Allele_56 | Res_5 | |
| H133 | Allele_105 | Res_4 | 1 |
| H134 | Allele_56 | Res_5 | |
| H135 | Allele_56 | Res_3 | |
| H136 | Allele_56 | Res_4 | |
| H143 | Allele_105 | Res_2 | |
| H145 | Allele_56 | Res_1 | |
| H146 | Allele_105 | Res_1 | |
| H147 | Allele_56 | Res_5 | |
| H148 | Allele_56 | Res_4 | |
| H151 | Allele_105 | Res_0 | |
| H152 | Allele_105 | Res_1 | |
| H153 | Allele_105 | Res_3 | |
| H156 | Allele_56 | Res_0 | |
| H161 | Allele_105 | Res_2 | |
| H163 | Allele_56 | Res_4 | |
| H167 | Allele_105 | Res_0 | |
| H176 | Allele_56 | Res_4 | |
| H177 | Allele_105 | Res_3 | |
| H178 | Allele_56 | Res_3 | |
| H179 | Allele_105 | Res_1 | |
| H181 | Allele_105 | Res_5 | 1 |
| H184 | Allele_105 | Res_2 | |
| H187 | Allele_56 | Res_3 | |
| H190 | Allele_56 | Res_3 | |
| H191 | Allele_105 | Res_2 | |
| H192 | Allele_105 | Res_1 | |
| H193 | Allele_105 | Res_2 | |
| H195 | Allele_105 | Res_3 | |

| | | | |
|---|---|---|---|
| H196 | Allele_105 | Res_1 | |
| H207 | Allele_105 | Res_0 | |
| H208 | Allele_56 | Res_3 | |
| H210 | Allele_56 | Res_2 | |
| H211 | Allele_105 | Res_0 | |
| H214 | Allele_105 | Res_3 | |
| H216 | Allele_105 | Res_4 | 1 |
| H217 | Allele_105 | Res_2 | |
| H223 | Allele_105 | Res_0 | |
| H225 | Allele_56 | Res_2 | |
| H226 | Allele_105 | Res_2 | |
| H227 | Allele_56 | Res_1 | |
| H228 | Allele_105 | Res_1 | |
| H230 | Allele_56 | Res_2 | |
| H231 | Allele_56 | Res_0 | |
| H232 | Allele_56 | Res_2 | |
| H234 | Allele_105 | Res_2 | |
| H235 | Allele_56 | Res_5 | |
| H236 | Allele_105 | Res_2 | |
| H237 | Allele_56 | Res_2 | |
| H238 | Allele_56 | Res_2 | |
| H242 | Allele_56 | Res_2 | |
| H244 | Allele_56 | Res_1 | |
| H247 | Allele_105 | Res_0 | |
| H248 | Allele_105 | Res_3 | |
| H251 | Allele_56 | Res_3 | |
| H252 | Allele_105 | Res_1 | |
| H255 | Allele_56 | Res_2 | |
| H256 | Allele_56 | Res_0 | |
| H258 | Allele_105 | Res_1 | |
| H259 | Allele_56 | Res_2 | |
| H263 | Allele_105 | Res_4 | 1 |
| H264 | Allele_56 | Res_3 | |
| H265 | Allele_105 | Res_4 | 1 |
| H266 | Allele_56 | Res_0 | |
| H267 | Allele_56 | Res_2 | |
| H272 | Allele_56 | Res_5 | |
| H274 | Allele_105 | Res_4 | 1 |
| H275 | Allele_56 | Res_2 | |
| H285 | Allele_105 | Res_4 | 1 |
| H286 | Allele_105 | Res_3 | |
| H287 | Allele_105 | Res_0 | |
| H289 | Allele_105 | Res_1 | |

| | | | |
|---|---|---|---|
| H291 | Allele_105 | Res_0 | |
| H295 | Allele_105 | Res_4 | 1 |
| H296 | Allele_105 | Res_3 | |
| H298 | Allele_56 | Res_1 | |
| H300 | Allele_105 | Res_4 | 1 |
| H304 | Allele_56 | Res_2 | |
| H305 | Allele_56 | Res_4 | |
| H306 | Allele_56 | Res_1 | |
| H307 | Allele_56 | Res_0 | |
| H308 | Allele_105 | Res_4 | 1 |
| H309 | Allele_105 | Res_1 | |
| H312 | Allele_56 | Res_1 | |
| H314 | Allele_56 | Res_1 | |
| H316 | Allele_56 | Res_3 | |
| H318 | Allele_56 | Res_4 | |
| H321 | Allele_105 | Res_0 | |
| H323 | Allele_105 | Res_4 | 1 |
| H324 | Allele_105 | Res_2 | |
| H327 | Allele_105 | Res_4 | 1 |
| H328 | Allele_56 | Res_0 | |
| H331 | Allele_56 | Res_3 | |
| H333 | Allele_56 | Res_0 | |
| H336 | Allele_56 | Res_1 | |
| H339 | Allele_105 | Res_2 | |
| H341 | Allele_56 | Res_1 | |
| H342 | Allele_105 | Res_3 | |
| H343 | Allele_56 | Res_2 | |
| H344 | Allele_105 | Res_3 | |
| H345 | Allele_56 | Res_4 | |
| H346 | Allele_56 | Res_2 | |
| H348 | Allele_56 | Res_2 | |
| H349 | Allele_105 | Res_2 | |
| H352 | Allele_56 | Res_2 | |
| H353 | Allele_56 | Res_3 | |
| H359 | Allele_56 | Res_1 | |
| H361 | Allele_56 | Res_1 | |
| H362 | Allele_105 | Res_3 | |
| H363 | Allele_56 | Res_1 | |
| H364 | Allele_56 | Res_5 | |
| H366 | Allele_105 | Res_1 | |
| H369 | Allele_56 | Res_1 | |
| H370 | Allele_56 | Res_3 | |
| H374 | Allele_56 | Res_1 | |

| | | | |
|---|---|---|---|
| H379 | Allele_56 | Res_2 | |
| H384 | Allele_56 | Res_1 | |
| H385 | Allele_105 | Res_5 | 1 |
| H386 | Allele_56 | Res_1 | |
| H388 | Allele_56 | Res_1 | |
| H392 | Allele_105 | Res_2 | |
| H398 | Allele_56 | Res_4 | |
| H401 | Allele_56 | Res_3 | |
| H402 | Allele_56 | Res_3 | |
| H405 | Allele_56 | Res_2 | |
| H406 | Allele_56 | Res_4 | |
| H408 | Allele_105 | Res_3 | |
| H410 | Allele_56 | Res_2 | |
| H411 | Allele_56 | Res_4 | |
| H412 | Allele_105 | Res_0 | |
| H414 | Allele_105 | Res_3 | |
| H416 | Allele_105 | Res_2 | |
| H418 | Allele_105 | Res_3 | |
| H419 | Allele_105 | Res_4 | 1 |
| H420 | Allele_105 | Res_4 | 1 |
| H421 | Allele_56 | Res_4 | |
| H422 | Allele_105 | Res_0 | |
| H423 | Allele_105 | Res_0 | |
| H424 | Allele_105 | Res_4 | 1 |
| H428 | Allele_105 | Res_2 | |
| H429 | Allele_105 | Res_2 | |
| H430 | Allele_56 | Res_4 | |
| H433 | Allele_56 | Res_4 | |
| H434 | Allele_105 | Res_5 | 1 |
| H435 | Allele_56 | Res_3 | |
| H438 | Allele_56 | Res_2 | |
| H440 | Allele_105 | Res_2 | |
| H441 | Allele_105 | Res_1 | |
| H446 | Allele_56 | Res_1 | |
| H447 | Allele_56 | Res_1 | |
| H449 | Allele_56 | Res_5 | |
| H450 | Allele_56 | Res_2 | |
| H451 | Allele_56 | Res_1 | |
| H452 | Allele_105 | Res_2 | |
| H455 | Allele_105 | Res_2 | |
| H462 | Allele_56 | Res_3 | |
| H463 | Allele_56 | Res_4 | |
| H464 | Allele_56 | Res_5 | |

| | | | |
|---|---|---|---|
| H466 | Allele_105 | Res_4 | 1 |
| H467 | Allele_56 | Res_5 | |
| H475 | Allele_105 | Res_5 | 1 |
| H477 | Allele_56 | Res_2 | |
| H478 | Allele_56 | Res_5 | |
| H482 | Allele_56 | Res_1 | |
| H484 | Allele_105 | Res_2 | |
| H485 | Allele_105 | Res_0 | |
| H486 | Allele_105 | Res_0 | |
| H489 | Allele_105 | Res_5 | 1 |
| H491 | Allele_105 | Res_1 | |
| H492 | Allele_56 | Res_2 | |
| H493 | Allele_105 | Res_0 | |
| H494 | Allele_105 | Res_1 | |
| H498 | Allele_105 | Res_2 | |
| H504 | Allele_56 | Res_0 | |
| H506 | Allele_56 | Res_0 | |
| H507 | Allele_56 | Res_2 | |
| H509 | Allele_105 | Res_0 | |
| H511 | Allele_56 | Res_1 | |
| H514 | Allele_105 | Res_3 | |
| H516 | Allele_105 | Res_2 | |
| H517 | Allele_56 | Res_3 | |
| H520 | Allele_105 | Res_1 | |
| H521 | Allele_56 | Res_3 | |
| H526 | Allele_105 | Res_1 | |
| H527 | Allele_105 | Res_0 | |
| H529 | Allele_105 | Res_2 | |
| H530 | Allele_56 | Res_4 | |
